# Aggregate Deep Visual Proteomics reveals selective organization of α-synuclein inclusions

**DOI:** 10.64898/2026.10.30.763157

**Authors:** Marc Oeller, Sabine Vasconez Grunauer, Cole S Sitron, Anne-Laure Mahul Mellier, Lucas Diedrich, Anton Schüle, Viktoria Ruf, Denys Oliinyk, Fabien Kuttler, Ulrich Dransfeld, Jana Prassler, Shani Ben-Moshe, Edwin Rodriguez, Tim Heymann, Enes Ugur, Katherine Madden, Yaoting Sun, Christina Ulrich, Yllza Jasiqi, Hilal A. Lashuel, F. Ulrich Hartl, Matthias Mann

## Abstract

α-Synuclein inclusions, the lesions typical of Parkinson’s disease, have been interpreted as protective deposits, toxic species or failed degradation products, each predicting a different composition and correlating that enrichment with neuron dysfunction and eventual degeneration. Distinguishing them requires measuring what an inclusion concentrates relative to the cell that built it. Here we developed Aggregate Deep Visual Proteomics, which uses ultrasensitive mass spectrometry to profile laser-microdissected inclusions against the remainder of the same cell or a matched host. In HEK293 cells, iPSC-derived dopaminergic neurons and post-mortem patient brain, each inclusion concentrated a selective subset of its cell’s proteome, independent of abundance and dominated by membrane, in neurons largely presynaptic. Inclusions took up whole assemblies, with subunits moving together in 421 of 438 complexes, and the V-ATPase alone was enriched in all three systems. Degradation cargo stayed with the receptors that route it, consistent with sequestration and failed degradation acting together. Neuritic and somatic inclusions differed, the former rich in UBQLN2 and the latter in intermediate filaments. In the cell line, phosphorylated α-synuclein was tenfold higher in the inclusion than in the rest of the cell, with about one third of the rise already present before an inclusion was visible. Measured against its host, the composition of an inclusion becomes a readout of how it formed and a source of candidate markers and therapeutic targets.

## Background

Pathological protein aggregation defines the major neurodegenerative diseases, with α-synuclein in Parkinson’s disease (PD)^1^, amyloid-β and tau in Alzheimer’s disease (AD)^2–6^, and TDP-43 in frontotemporal dementia (FTD)^7,8^. In PD, the fastest-growing^9^ and second most common^10^ of these disorders, α-synuclein is the principal but not the sole component of Lewy bodies (LB) and Lewy neurites (LN)^1^. How LBs and LNs contribute to disease progression remains unclear. Models have cast LBs as protective sequestration sites for toxic aggregates and damaged organelles, as toxic species themselves, or as failed degradation products. Each role implies a different aggregate composition, and none has been mapped against the cell the inclusion formed in.

Lewy pathology is heterogeneous, differing by morphology, brain region, disease stage and patient, and progression modeling resolves at least three distinct trajectories^11,12^. Heterogeneity extends to the subcellular scale, where one neuron can carry morphologically distinct neuritic and somatic inclusions. Imaging has begun to expose their internal structure. Electron and correlative light-electron microscopy reveal inclusions crowded with membranes and organelles^13,14^, and STED microscopy shows α-synuclein proteoforms spatially organized within them^15^. These differences may track which complexes and organelles each inclusion sequesters. These methods remain limited to a few markers, short of a proteome-level description. Whether these forms are stages of one process or products of distinct pathways is unresolved, and the answer bears on which drives neuronal dysfunction.

The first constituents identified were histopathological markers, among them ubiquitin^16^, p62/SQSTM1^17^, neurofilaments^18^, and phosphorylated α-synuclein. Phosphorylation at Ser129 (pS129) is the most consistent, carried by about 90% of the α-synuclein deposited in LBs against about 4% of the soluble protein in normal brain^19,20^. They identify an inclusion as pathological but separate no form or stage from another. Insoluble-fraction biochemistry cataloged the aggregated proteome^21–24^, limited-proteolysis mass spectrometry (MS) resolved conformational states^25^, proximity labeling mapped the protein neighborhood of phospho-α-synuclein (∼261 proteins in human brain)^26^, and laser-capture microdissection recovered the proteomes of cortical inclusions^27^. These inventories could not determine whether a protein is concentrated in the inclusion, belongs to the surrounding cell or is induced by stress. They also average over inclusions of different morphology and location, so neuritic and somatic forms have never been compared directly. The open question is what an inclusion concentrates relative to its host, and whether that differs between the forms Lewy pathology takes.

Spatial proteomics^28–32^ now yields deep proteomes from minute tissue areas by coupling laser microdissection to ultrasensitive mass spectrometry^33–36^, and Deep Visual Proteomics (DVP) brought this to cell-type resolution *in situ*, linking AI-guided image analysis to automated laser microdissection^37–40^. In AD brain, profiling tangle-bearing neurons against spared neighbors resolved the cell carrying the pathology but left the aggregate diluted into the whole-cell proteome^41^.

Here we developed Aggregate DVP, which brings DVP to the subcellular scale. It isolates morphologically defined inclusions by automated segmentation and microdissection, and profiles them against a matched host fraction from the same sample, in a cell line expressing A53T α-synuclein, in iPSC-derived dopaminergic (iDA) neurons^42^ and in post-mortem brain from patients with LB disease. This let us ask whether an inclusion is a selected subset of its host cell or the random end-state of aggregation, whether selection acts on single proteins or on assemblies, and whether neuritic and somatic inclusions differ. It proved to be a selected subset, with selection acting on complexes and organelles, varying with the compartment it forms in. It extended to the pS129 that defines the pathology but not to the enzymes that add and remove it, about one third of it in place before an inclusion was visible. Composition answers of this kind separate the three interpretations. They also put within reach what makes inclusions differ, in what order their contents arrive, and which losses the host cell cannot compensate.

## Results

### Aggregate DVP measures each inclusion against the cell that formed it

A protein found in an α-synuclein inclusion is not necessarily concentrated there and may only reflect the composition of the surrounding cell. Separating the two requires a comparator from the same cell, which an inclusion does not readily provide. It has no fixed location or boundary, is defined only by its morphology and is smaller than the cell that contains it. In Aggregate DVP, we isolate the inclusion itself and profile it against a host fraction from the same sample, either immunostained cells or archival chromogenic tissue. In HEK293 cells, the host is what remains of that same cell once the inclusion has been cut out. In iDA neurons it is a matched soma for a somatic inclusion and a matched neurite for a neuritic one, and in brain a matched DAB-positive neuron, each cut from the same slide. Every enrichment value is therefore a ratio against matched host material from the same sample. Because an inclusion and its host share fixation, batch and donor, these variables are constant within each comparison.

Across all three systems, the workflow is shared (Figure 1a): pathological inclusions are stained and imaged, computationally segmented, excised by laser microdissection, and profiled by low-input MS. All samples were stained for pS129, the standard marker of pathological α-synuclein inclusions^19,20,43^, and imaged by high-content or slide-scanning microscopy, with fluorescent counterstains for the cell models, and 3,3ʹ-diaminobenzidine (DAB) immunohistochemistry with cresyl violet for post-mortem brain, the standard modality in clinical neuropathology (Figure 1b, S1a-c).

**Figure 1.**
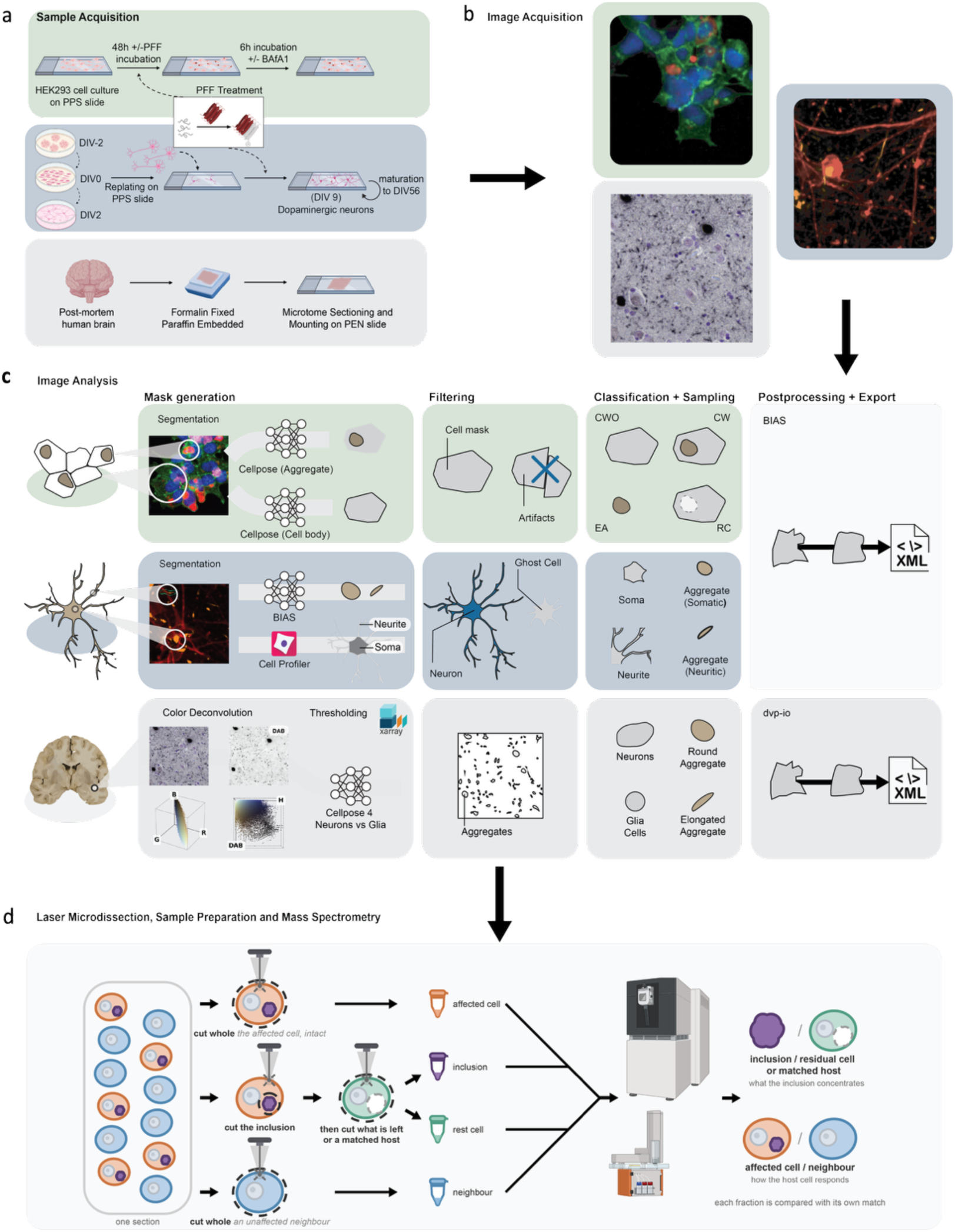
The Aggregate DVP workflow across three systems of α-synuclein aggregation. (a) Sample preparation for the three systems: HEK293 cells stably expressing A53T α-synuclein, ± pre-formed fibrils (PFFs) and ± Bafilomycin A1 (BafA1); iDA neurons at DIV56 ± PFFs; and formalin-fixed, paraffin-embedded (FFPE) post-mortem brain on membrane slides. (b) Staining and representative imaging (more detailed images in Figure S1): all systems stained for pS129 with a cellular or tissue counterstain. (c) Image analysis defining the laser-microdissectable fractions per system. HEK293: cells with aggregates (CW), cells without aggregates (CWO), excised aggregates (EA), and the residual cell body after aggregate excision (RC). iDA neurons: somatic and neuritic aggregates and their host fractions. Brain: pS129-positive aggregates classified as round (Lewy body-like) or elongated (Lewy neurite-like), with host cell bodies classified as neurons or glia. (d) Laser microdissection and proteomic readout. From the same sample, four fractions are collected: whole cells carrying an inclusion (CW); the inclusion itself (EA), excised first; the remainder of the same cell, excised second (residual cell body, RC), or a matched host fraction; and whole cells without an inclusion (CWO). Fractions are processed by automated low-input sample preparation and analyzed by LC-MS/MS. Two ratios are computed. EA/RC, within the same cell, gives what the inclusion concentrates relative to its own host. CW/CWO, between cells, gives the response of a cell to carrying an inclusion; only the first isolates the inclusion itself.

Segmentation produced polygons marking the regions to excise (Figure 1c), and what it had to resolve differed by system. In HEK293 cells, custom segmentation^44^ models separated aggregates from cell bodies, and overlaying the masks gave four fractions, the excised aggregate (EA), the residual cell body it was cut from (RC), and whole cells with and without an inclusion (CW, CWO). These support two comparisons, EA against RC for what an inclusion concentrates relative to the rest of its own cell, and CW against CWO for what changes in a cell that carries one. In iDA neurons and in brain, each inclusion was cut against a host fraction from its own compartment, somatic inclusions against a matched soma and neuritic ones against a matched neurite, round brain inclusions against a DAB-positive neuron and elongated ones against the surrounding neuropil, with glia as a further fraction. Enrichment is therefore read against the compartment an inclusion formed in and not against a neuronal average. All fractions were located automatically, whereas prior immunohistochemistry-based proteomics had relied on manually selected regions^45–47^.

Compact inclusions span roughly 50 to 100 µm² in cross-section, well above the ∼1 µm precision of laser microdissection, so each is excised whole, at up to ∼1,000 shapes per hour verified by imaging before and after dissection, with tens of thousands of shapes cut across the three systems (Figure 1d, Figure S1d). Thin neuritic inclusions (∼20 µm²) approach this limit in their narrow dimension and were cut with a surrounding margin to recover them whole. The three systems differ in what they reproduce. A cell line allows aggregation to be induced and manipulated, iDA neurons place Lewy-like inclusions in their own compartments, and patient brain supplies the pathology itself, so agreement across all three is the strongest evidence a finding is general.

### HEK293 inclusions are selective compartments tuned by clearance state

To test whether an inclusion is a distinct compartment, we began with HEK293 cells stably expressing A53T α-synuclein and seeded with pre-formed fibrils (PFFs). We profiled four fractions (CW, CWO, EA, RC; Figure 1c) in six wells per condition including PBS-treated controls, and quantified 7,290 proteins (Figure 2a, Figure S2a, b). Fibril exposure alone changed the host cell before any inclusion was visible. In cells without visible aggregates, 371 proteins increased and 225 decreased relative to PBS-treated controls (n=6, Figure 2b), α-synuclein among the strongest hits, protein degradation and translation up, and mitochondrial respiration, membrane traffic and solute transport down (Figure S2c). Each inclusion is measured against the remainder of a cell in this altered state.

**Figure 2.**
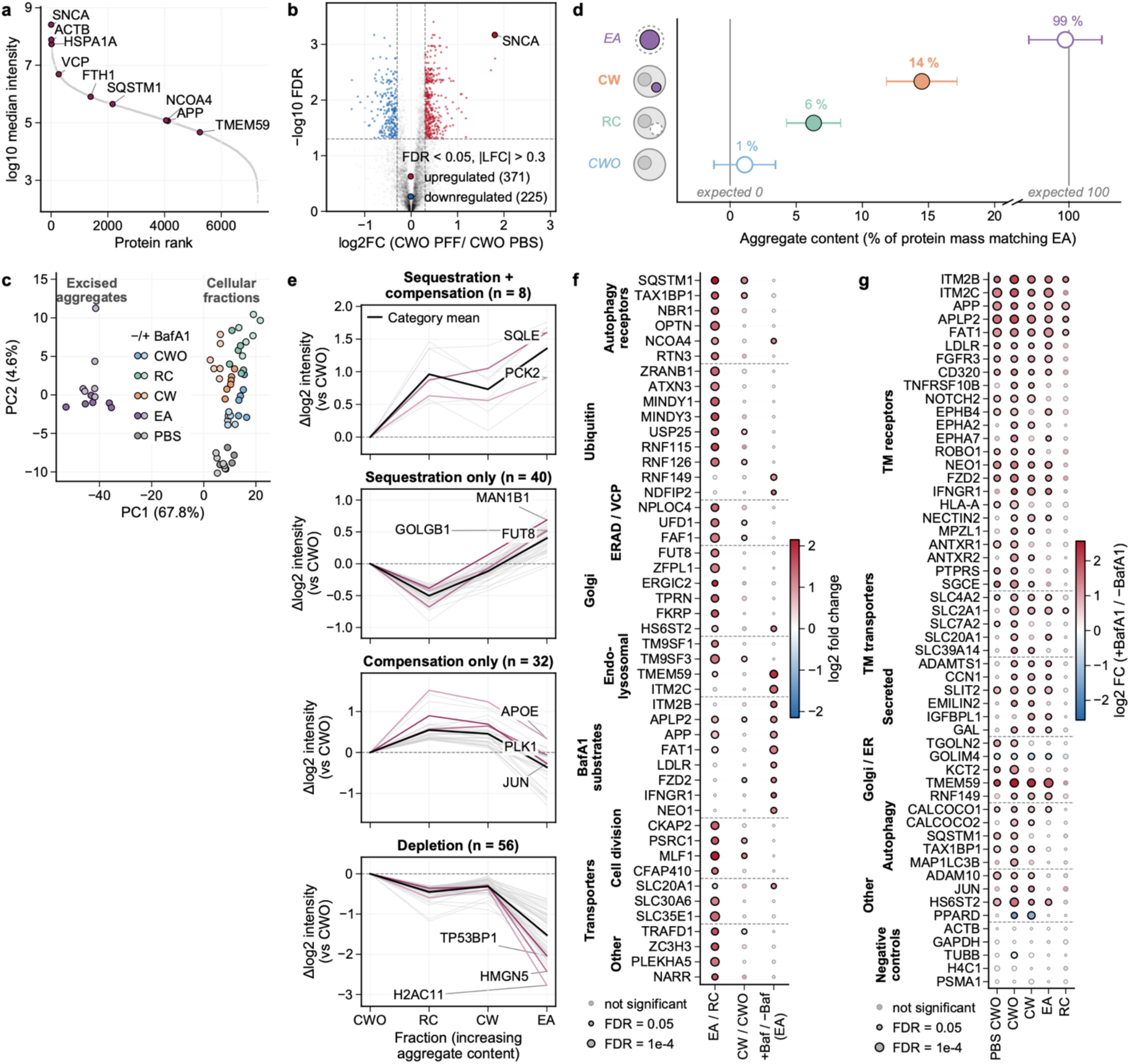
Aggregate DVP of HEK293 cells expressing A53T α-synuclein, with and without lysosomal blockade. (a) Rank plot of median protein-group intensity across all fractions (n=6, 7,290 proteins quantified); selected proteins labeled. (b) Volcano plot of PFF-versus PBS-treated cells without aggregates (−BafA1). (c) Principal component analysis (PCA) of all fractions. (d) Aggregate content of each dissected fraction, as the percentage of its protein content matching the excised aggregate (−BafA1; CWO defines 0%, EA defines 100%). Positions are the median of per-protein estimates across 1,124 proteins differing at least twofold between the two reference fractions (BH-FDR < 0.05). Reference fractions are placed by leave-one-out (CWO 1.1%, EA 99.3%). Points, mean of six wells; error bars, SD. (e) Trajectory analysis across CWO, RC, CW, and EA (−BafA1), showing four protein-redistribution patterns with selected proteins labeled per pattern. y-axis ranges differ between panels; the four patterns are classes of the ANOVA and post-hoc tests (Methods), and magnitudes are not comparable across panels. Gray lines, individual proteins; bold black line, pattern mean; colored lines, labeled examples. (f) Dot plot of the most aggregate-enriched proteins (Methods), grouped by biological category. Columns: the excised aggregate against its matched host fraction (EA / RC) and cells with against cells without aggregates (CW / CWO), both −BafA1, and the BafA1 effect within the aggregate. Dot size encodes −log10(FDR), color encodes log2 fold change. (g) Dot plot of BafA1-responsive proteins across five fraction comparisons, grouped by biological category; dot size encodes −log10(FDR), color encodes log2 fold change.

Capture was not driven by abundance. Instead, the aggregate separated from every cellular fraction along the dominant axis of variation (68% of variance; Figure 2c). Its enrichment was independent of a protein’s cellular level (r = −0.16; Figure S2d), and it was enriched in ER, Golgi and mitochondrial proteins and depleted of nuclear components (Figure S2e). The two dissected parts should also sum to the intact cell. Aggregate-bearing cells show 14.5 ± 2.7% aggregate content and the residual cell body 6.3 ± 2.0% (Figure 2d, Methods), so the excised region holds about a seventh of the cell’s protein. The two parts reconstructed the intact cell to within 1.4 times measurement noise, and every alternative model fit worse (1.6 to 42.7 times; Figure S2f). The inclusion is therefore a discrete, selective compartment.

Pairing each aggregate with its host also shows whether the cell compensates for proteins drawn into the inclusion (Figure 2e, Methods). Of the proteins enriched in the aggregate, 40, mostly Golgi and glycosylation machinery, were depleted from the host, while only 8, including the cholesterol-synthesis enzyme SQLE, were increased. The 40 are consistent with incidental capture without replacement, while the 8 are consistent with cargo marked for degradation. The inclusion therefore draws on protein the cell already has and replaces little of it, a potential loss of function.

The proteins retained most strongly against the residual cell body fell into four classes, selective autophagy receptors including ferritinophagy receptor NCOA4, an inclusion-resident ubiquitin-editing machinery, the VCP/p97 segregase and its cofactors, and Golgi and endolysosomal membrane proteins (Figure 2f). Gene-set enrichment ranked ER glycosylation and ER-to-Golgi transport highest, followed by the respiratory chain, ER membrane insertion, Golgi and endosomal transport and ER-associated degradation, while nuclear, chromatin and RNA-processing categories were depleted (Figure S3a). The inclusion therefore holds clearance machinery alongside substrates of that machinery. Most were significant only against the residual cell body, so a comparison of cells with and without an inclusion would have missed them.

To test whether composition tracks the clearance state of the cell, we profiled the same fractions with and without the lysosomal inhibitor Bafilomycin A1 (BafA1)^48,49^. This gave the lysosome-dependent proteins of each fraction (Figure S3b, Methods). The response was selective, with canonical autophagy markers, aggrephagy receptors and NCOA4 accumulating upon inhibition, while mitophagy sensors did not. Aggrephagy receptors accumulated more in fibril-exposed cells without an inclusion than in the host cell body of cells with one. The responding subset differed from the proteins that responded cell-wide, so the effect is specific to the inclusion. These proteins were almost all membrane-associated (Figure 2g). The categories that define the inclusion at baseline stayed enriched (Figures S3a, c, d), so clearance state tunes what the inclusion holds without rebuilding it. One responding receptor ties the inclusion to a recognized feature of PD, iron dyshomeostasis^50^. The ferritinophagy receptor NCOA4 was co-sequestered with its cargo (FTL/FTH), confirmed by anti-FTL immunofluorescence (Figure S3e, f). Inclusion formation thus engages the iron-handling machinery implicated in dopaminergic oxidative stress^51,52^.

The HEK293 inclusion is therefore a compartment of its cell, built around autophagy receptors, VCP cofactors and organellar membranes, tuned by clearance state and assembled largely from protein the cell does not replace.

### Neuritic and somatic α-synuclein inclusions exhibit distinct proteomic signatures

Neuritic and somatic inclusions coexist within single neurons, and whether they are successive stages of one process or products of distinct aggregation pathways is unresolved. We profiled both types in human iDA neurons seeded with PFFs, a model that develops pathology without α-synuclein overexpression, reproduces the inclusion types seen in patients, and in which neuritic aggregates mature into membrane-crowded, LB-like somatic inclusions^42^. Inclusions were selected on the published criteria of pS129 and p62 staining with morphology. From a single culture we then microdissected somata and neurites separately, collecting inclusion-free regions alongside neuritic (NAgg) and somatic (SAgg) inclusions, and quantified 6,444 proteins across 33 samples (Figure 3a, Figure S4a, b). PCA separated soma from neurites and PFF-from PBS-treated samples (Figure 3b), and 35 of 36 soma and 24 of 44 neurite markers were enriched in the expected direction in controls (Figure 3c, Figure S4c).

**Figure 3.**
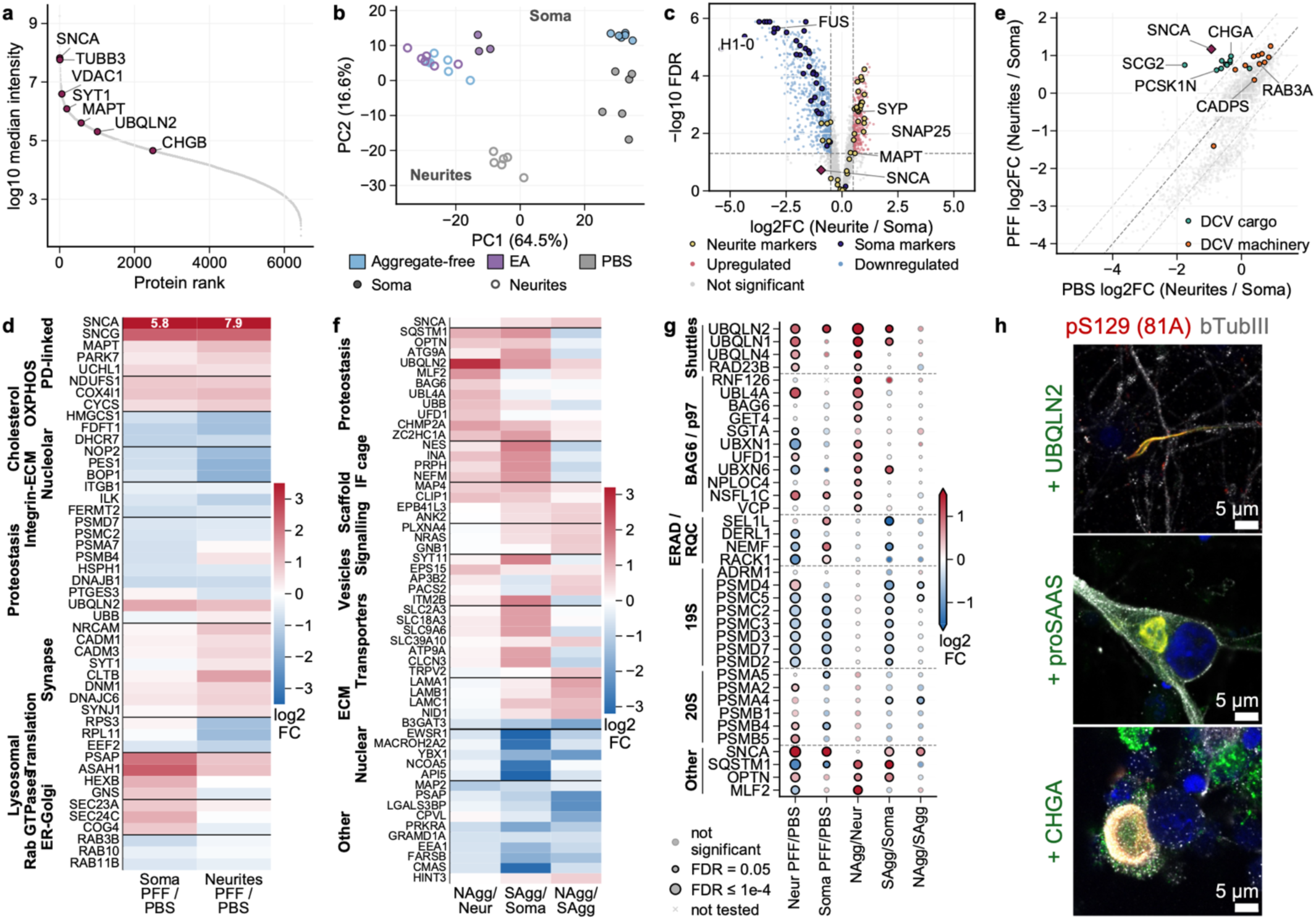
Aggregate DVP of neuritic and somatic α-synuclein inclusions and their host fractions in iDA neurons. (a) Protein rank-abundance plot across all iDA neuron samples, ordered by median log2 intensity; disease-relevant and compartment-marker proteins labeled. (b) PCA of all 33 samples across six compartment-condition groups (PBS / PFF / aggregate × soma / neurites; n = 6 per group, except PFF somatic aggregates, n = 3). (c) Volcano plot of PBS-treated neurites versus PBS-treated soma; established soma (dark blue) and neurite (yellow) markers and α-synuclein labeled. (d) Log2 fold-change heatmap of the compartment-specific PFF response (PFF versus PBS) in soma (left) and neurites (right); proteins grouped into functional modules, with off-scale SNCA fold-changes annotated in the panel. (e) Two-dimensional scatter of the dense-core vesicle (DCV) proteome: x-axis, log2 fold change neurites/soma in PBS; y-axis, the same in PFF. DCV cargo, teal; DCV machinery, orange. Dashed grey line, identity (y = x); dashed light grey lines, identity ± 1 log2 unit. (f) Log2 fold-change heatmap of aggregate composition: neuritic aggregates versus PFF neurites (left), somatic aggregates versus PFF soma (middle), and neuritic versus somatic aggregates (right); proteins grouped into functional modules. (g) Dot plot of ubiquitin-dependent quality-control proteins across the five contrasts; color, log2 fold change; dot size, FDR; rows grouped by function, with the 19S and 20S rows showing the ubiquitin receptors PSMD4 and ADRM1 plus three representative subunits per subcomplex; computed on the 6,372 protein groups left after contaminant removal, Welch t test with Benjamini-Hochberg correction over all proteins tested in each contrast. (h) Immunofluorescence of PFF-seeded iDA neurons stained for UBQLN2 in inclusion-bearing neurites (top), proSAAS/PCSK1N (middle) and chromogranin A (bottom) in inclusion-bearing soma; target green, pS129 α-synuclein red, nuclei blue, b-tubulin III grey. Scale bars, 5 µm. Channels separately shown in FigureS5.

Fibril exposure changed soma and neurites differently even where no aggregate was visible (Figure 3d, Figure S4d). Translation machinery increased in soma and fell in neurites, and tau increased only in neurites. The proteasome fell in soma, while in neurites the 19S particle fell and the 20S core rose. α-Synuclein rose 54-fold in soma and 234-fold in neurites, values that include internalized fibrils. Each inclusion type therefore forms in a differently altered environment, though inclusion-free regions may hold aggregates below detection.

Aggregate DVP separates redistribution within the cell from capture by the inclusion, and dense-core vesicles show the two running in opposite directions (Figure 3e). Following PFF exposure, granins, VGF and prohormone-processing enzymes were relatively enriched in neurites without visible inclusions, yet were selectively concentrated in somatic inclusions (Figure S4e), so capture ran against local availability. Both held organellar material from mitochondria, lysosomes, ER and Golgi (Figure S4f).

Each measured against its own compartment, the two inclusion types accumulated different proteins. The neuritic inclusion was enriched for machinery that handles misfolded and ubiquitinated proteins, led by UBQLN2 (7-fold), together with endocytic and ESCRT proteins (Figure 3f). The somatic inclusion was enriched more strongly for intermediate filaments, which form the cage of aggresomes, and held more membrane transporters and less nuclear protein than its soma. Both contained more p62/SQSTM1 than their hosts, far more so in the somatic inclusion. Immunostaining detects p62 inconsistently in neuritic pathology, while mass spectrometry quantified it directly. α-Synuclein showed little enrichment over PFF-exposed neurites (+0.04 log2) or soma (+0.44), likely because both hosts already carried fibril-derived protein, so composition extends well beyond α-synuclein itself. These distinct signatures may reflect successive maturation stages or compartment-specific pathways of inclusion formation.

UBQLN2, which shuttles ubiquitinated substrates to the proteasome, showed the largest enrichment in the neuritic inclusion, so we examined the wider ubiquitin-dependent quality-control machinery in both inclusions and their hosts (Figure 3g). The other ubiquilins increased, as did BAG6 with its ligase RNF126 and the p97 extraction machinery, which together recognize exposed hydrophobic segments, ubiquitinate them and extract them from aggregates. Proteasomal receptors, ERAD and ribosome-associated quality control stayed at host level, so recognition and extraction machinery was recruited while downstream degradation components were not. The somatic inclusion shared only the ubiquilins and RNF126. Staining confirmed this, with UBQLN2 bright in pS129-positive neuritic inclusions and proSAAS/PCSK1N and chromogranin A in somatic ones (Figure 3h, S5).

The two inclusion types therefore carry distinct compositional signatures, ubiquitin-dependent quality control in neurites and cytoskeletal and membrane proteins in somata.

### Round Lewy inclusions in patient brain are membrane-crowded, reproducibly across donors and biobanks

Neither cell model reproduces how clinical inclusions begin, since both start from added fibrils that may differ biochemically and structurally from those formed in patients. We therefore applied Aggregate DVP to formalin-fixed post-mortem tissue from donors with PD and Lewy body disease and from neurologically healthy controls, drawn from two independent biobanks. Sections were immunostained for pS129 α-synuclein with DAB and counterstained with cresyl violet, cell bodies were classified as neurons or glia, and inclusions as round (Lewy body-like) or elongated (Lewy neurite-like) on the DAB signal, size and overall shape. These two classes were not subdivided further by morphology. Neuropil was cut as randomly placed circles in regions free of cell bodies and inclusions. Across five disease donors and five controls, we analyzed 332 samples across seven fractions and quantified 6,842 protein groups (Figure 4a, Figure S6a, b).

**Figure 4.**
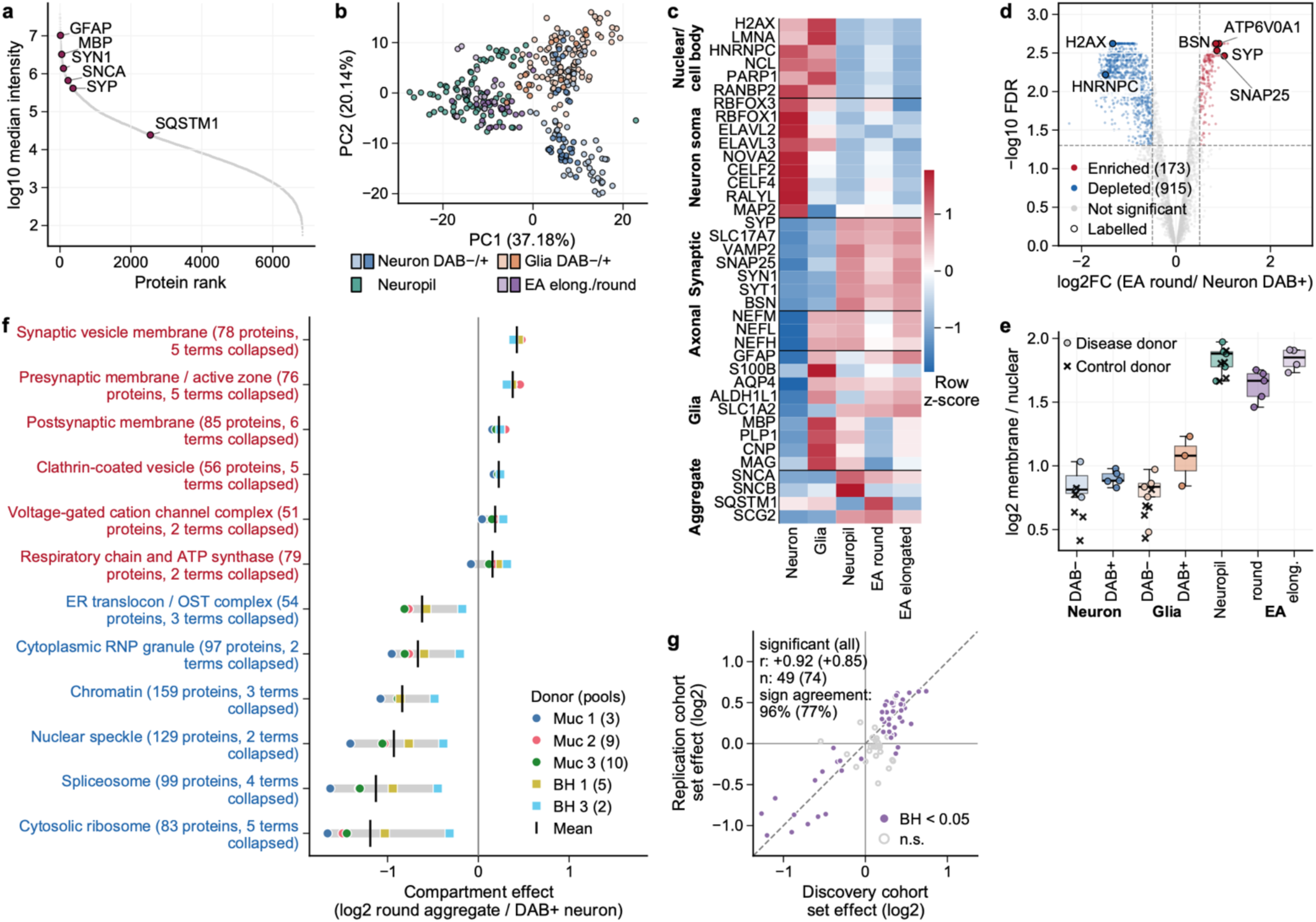
Aggregate DVP of post-mortem human brain tissue across two independent biobanks. (a) Rank-abundance plot of median protein-group intensity; selected proteins labeled. (b) PCA of all 332 samples, colored by fraction (neuron DAB−/DAB+, glia DAB−/DAB+, neuropil, round aggregate, elongated aggregate; 13 to 96 samples per fraction). PC1 (37%) separates cell bodies from neuropil and aggregates; PC2 (20%) separates glia from neurons. (c) Marker composition across five cell types (neurons, glia, neuropil, round and elongated aggregates); color, per-marker z-score of log2 intensity, rows grouped by cell class and compartment. (d) Volcano plot of round aggregates versus their DAB-positive host neurons (n = 5 paired donors, eBayes moderated t on donor means). Dashed lines, |log2FC| > 0.5 and FDR < 0.05; 173 proteins aggregate-enriched and 915 higher in the host soma. (e) Per-fraction log2 ratio of summed membrane/vesicular to soluble nuclear/translation protein intensity. Colored dots, disease donors; gray crosses, controls; boxes, median and interquartile range (IQR); whiskers, 1.5 × IQR. (f) Donor-level log2 ratio of round aggregate to matched host DAB+ neuron for GO cellular component terms, the six most enriched and six most depleted after collapsing overlapping terms (FDR < 0.05, gene set enrichment on the region-paired fit, Methods). Colored dots, individual donors; circles, Munich; squares, Banner; gray bars, range across donors; black ticks, mean; parentheses, detected proteins and number of terms collapsed. (g) Term-level values (mean of the detected members minus that of all other proteins) for GO cellular component terms, discovery against replication cohort, with significance from the discovery cohort only (one-sample t test, Benjamini-Hochberg). Filled dots, FDR < 0.05; open dots, not significant; dashed line, identity; parentheses, the same statistics over all tested terms.

The dominant axes of variation were biological, with no clustering by plate, sex, donor, brain bank or region (Figure 4b, Figure S6c–f). Canonical markers confirmed each fraction, with p62/SQSTM1 confined to round inclusions (Figure 4c). These inclusions retained the regional profile of their host neurons (Figure S6g), as expected if they form from their host’s material.

Compared with the neuron that contained it, a round inclusion held a distinct set of proteins (Figure 4d). It was enriched in synaptic-vesicle and SNARE proteins, the active-zone scaffold bassoon and the V-ATPase subunit ATP6V0A1 and depleted of nuclear and RNA-processing machinery, in all five paired disease donors. The aggregate does not simply report the state of a diseased neuron, because no protein differed significantly between aggregate-free neurons from disease and control donors, and the aggregate-versus-host and disease-versus-control signatures share only about a third of their variance (Figure S6h, i, Methods).

The ratio of membrane and vesicle protein to soluble nuclear and translation protein was 1.6-fold higher in round inclusions than in the cell bodies around them, across cell types and donors (Figure 4e, Figure S6j, Methods). This measures, as a protein ratio, the membrane and organelle crowding that structural studies have described^13^. Elongated inclusions were close to the surrounding neuropil (Figure 4e), so at this resolution only round inclusions resolve as a discrete structure. Thin, neuropil-embedded inclusions may still have a distinct proteome that surrounding tissue masks.

With donor as the statistical unit, the compartments enriched in round inclusions over their matched DAB-positive host neurons were the presynapse, clathrin-coated vesicles, cation channels and the respiratory chain, and the depleted ones were ribosomes, spliceosome, chromatin and RNP granules (Figure 4f). Direction was shared, with 165 of the 181 significant terms moving the same way in all five donors relative to the rest of the proteome. Magnitude was shared for the enriched compartments, whereas the two brain banks differed in the magnitude of depletion, a difference that cannot be separated from differences in post-mortem interval and measurement depth. Patient-specific variation was bounded at about 10% of the variance between pools (Figure S6k, l, Methods).

Three further PD cases with three matched controls, profiled as an independent batch and searched jointly with the discovery cohort (443 samples), reproduced the signature, with compartment-level effects correlating at r = +0.92 (96% directional agreement across the terms called in discovery) and protein-level effects at r = +0.88 (Figure 4g, Figure S6m). In every donor of both cohorts membrane and vesicle proteins made up more of the round inclusion than of its own host neuron (Figure 4e, Figure S6n).

Thus, in donors all at neocortical Lewy body stage, α-synuclein inclusions in patient brain are membrane- and organelle-crowded structures, enriched in synaptic-vesicle, SNARE and V-ATPase proteins relative to their host neurons.

### Across three systems, enrichment follows complex and organelle membership

We next compared each inclusion with its host across the three systems, to separate general features from system-specific ones and to ask how closely the models reproduce authentic Lewy pathology. Across the 2,034 proteins quantified in all three, every pairwise correlation was positive, closest between the two neuronal systems (r = 0.80) and more modest for HEK293 (r = 0.4 to 0.5; Figure 5a). Seven proteins were enriched in the inclusion and 256 depleted in all three systems. Four belong to the V-ATPase, enriched in its entirety (Figure S7a), two are secretory-vesicle membrane proteins (VAMP2, SV2A) and the seventh is GGT7, a membrane glutathione hydrolase not previously linked to α-synuclein, so all seven are membrane-associated. Dense-core vesicles show the same selectivity, their cargo and release machinery retained while the biogenesis factor HID1 is excluded (Figure S7b), so parts of one functional unit are not necessarily taken up equally.

**Figure 5.**
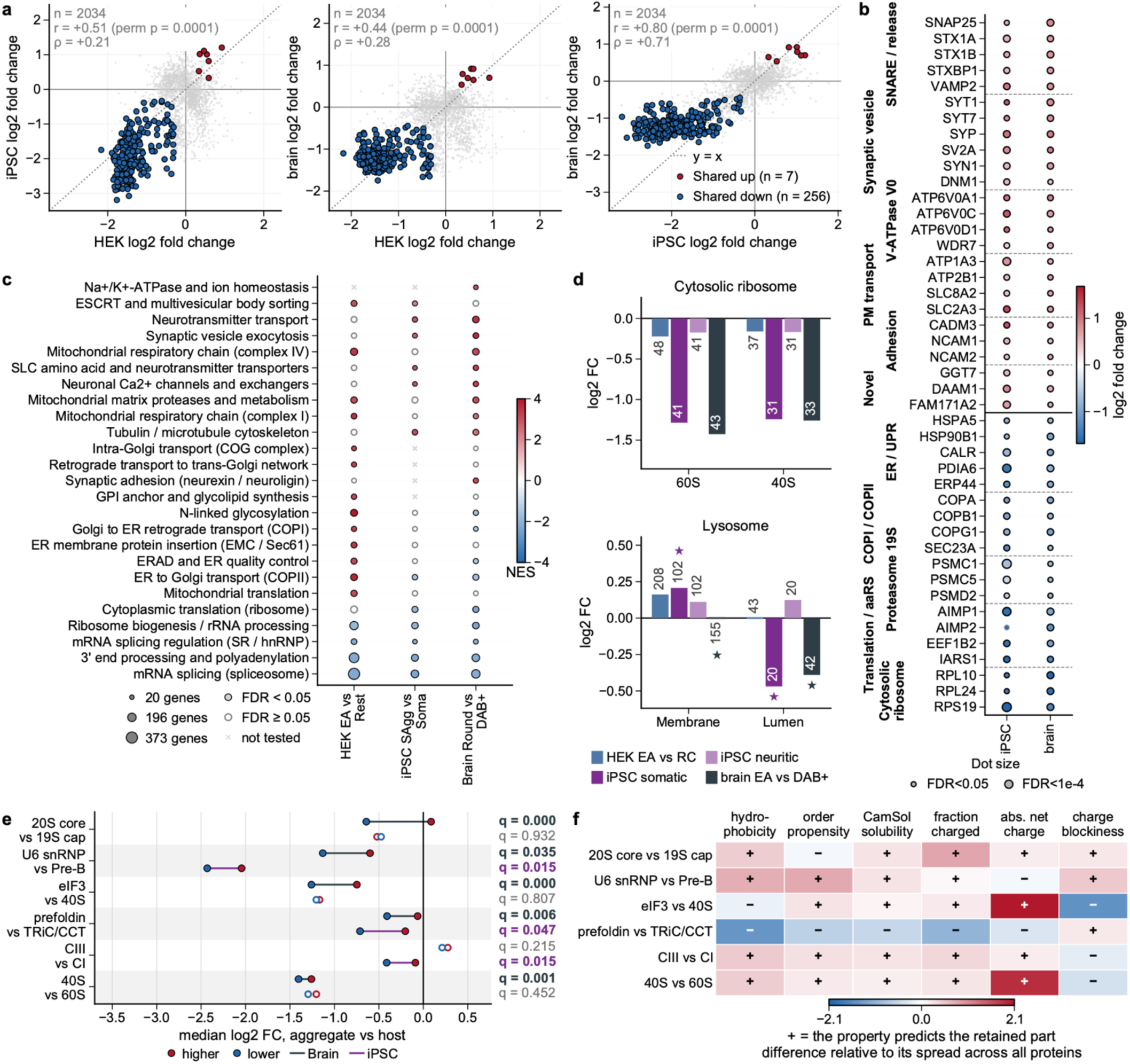
A shared α-synuclein inclusion signature across three systems, with complexes and organelle modules mostly enriched or depleted coherently. (a) Pairwise scatter of aggregate-versus-host log2 fold change for the 2,034 protein groups quantified in all three aggregate-versus-host contrasts (HEK EA vs RC, −BafA1; iDA SAgg vs PFF soma; brain round vs neuron DAB+). Red, shared enriched (n = 7); blue, shared depleted (n = 256); gray, other. Text in each panel gives Pearson r with its permutation p and Spearman ρ (n = 2,034); dashed line, y = x. (b) Selected proteins of the iDA and brain signature (47 enriched, 476 depleted; |log2FC| ≥ 0.3 and FDR < 0.05 in both, concordant sign), grouped by function. Dot color, log2FC; dot size, −log10 FDR. (c) Cross-system aggregate-versus-host GSEA (Reactome and GO) across three contrasts; rows, terms with FDR < 0.05 in at least one contrast, deduplicated per theme. Color, normalized enrichment score; size, detected set size; filled, FDR < 0.05; grey cross, not tested. (d) Module scores for one organelle excluded as a single piece and one that separates internally. Bars, group difference in per-sample module score per aggregate vs host contrast (brain, mean of the per-donor differences); numbers, member count; star, module in at least one separating pair (Methods). (e) Six pairs of directly interacting assembly parts that separate in at least one contrast; brain round vs DAB+, charcoal; iDA SAgg vs soma, purple. Dots, median log2FC of that part’s quantified members; red, the higher part; blue, the lower, q < 0.05 in that contrast; gray, not significant. Mann-Whitney between the two parts’ member log2FC, BH within contrast. (f) Sequence-property difference between the two parts of each pair in (e), oriented so that positive matches the proteome-wide prediction (higher part more hydrophobic and ordered, less soluble, charged and charge-blocky). Color, difference in SD units of that property across all quantified proteins; glyph, sign. Values derive from sequence and apply to both contrasts.

Requiring agreement only between the two neuronal systems gave a fuller picture (Figure 5b, Figure S7c). The enriched group is dominated by the presynaptic terminal, its release machinery and synaptic vesicles, together with the V0 sector of the V-ATPase, plasma-membrane transport and synaptic adhesion molecules, and the depleted group by ER chaperones, the COPI and COPII coats, the proteasome 19S particle and the translation machinery. ALS and FTD RNA-binding proteins were depleted more strongly than nuclear proteins overall (Figure S7d), so a protein’s own aggregation propensity does not predict enrichment.

Of the 228 proteins reported as aggregate constituents by three or more earlier studies (Methods), we detected 96% in brain, but only 25% were enriched relative to matched host neurons, mainly presynaptic and membrane proteins (Figure S7e, f). SYT7, DAAM1, NCAM2, GGT7 and FAM171A2^53^ had not appeared in earlier α-synuclein inclusion datasets (Figure 5b). Pathway enrichment separates the systems (Figure 5c), with the ER and Golgi set enriched in HEK293 alone, as expected if overexpressed α-synuclein binds secretory membranes, a load neurons at endogenous dose do not carry. Both neuronal systems instead share synaptic enrichment and depletion of translational machinery.

Enrichment followed structures more than single proteins. We divided 16 organelles into 46 modules and scored every complex with at least four quantified proteins per module (Figure 5d, e, Figure S8 a-m, Methods). Only 33 of 201 module pairs differed by more than 23%, and complex subunits were less dispersed than in matched random sets in 247 of 255 HEK, 72 of 74 iDA and 102 of 109 brain complexes. Most complexes were depleted relative to their host, 169, 53 and 70 significantly so against 33, 5 and 15 enriched. Where a structure split, the split ran along an internal boundary, the lysosomal membrane above its lumen in fold change and the mitoribosome below the rest of the mitochondrion. In five of the six complexes that separated, the higher part was itself still depleted, so these splits reflect differential loss. Carry-over of surrounding cytoplasm cannot produce such splits, because it would transfer interacting parts in their cellular proportions. These results are consistent with organellar material, whole or damaged, entering the inclusion with its internal proportions largely preserved, with defined exceptions^13,14^.

We then asked whether physical character predicts enrichment, from sequence-derived hydrophobicity and predicted solubility (CamSol^54,55^). In brain, hydrophobicity correlated with enrichment the more strongly for larger units, from ρ = +0.21 across proteins to +0.61 across the most and least enriched complexes, whereas no property exceeded ρ = +0.11 among the subunits of one complex (Figure S8n, Methods). Physical character therefore separates assemblies from one another but not their own subunits, which are usually enriched to a similar degree. Six complexes did split, and in five the part higher in the aggregate was the less soluble and less charged one. Prefoldin is the sixth, the more soluble and more charged partner of TRiC and yet the less depleted, so at least one determinant of retention is not physicochemical (Figure 5f). Members of a complex are therefore enriched or depleted together.

### Phosphorylated α-synuclein accumulates in the inclusion while its enzymes do not

Whether pS129 precedes the visible inclusion has remained unresolved. We extended Aggregate DVP to the phosphoproteome with nanoPhos, our ultrasensitive phospho-enrichment workflow^56^. In our HEK293 model we normalized each phosphosite to its parent protein, measured from the same material, so a rise reflects modification and not more protein. Across 4,622 localized (Class-I) phosphosites in 41 samples, three to five per fraction and arm, the excised aggregates separated from every cellular fraction along an axis that tracks α-synuclein phosphorylation (Figure 6a, b, Figures S9a-e).

**Figure 6.**
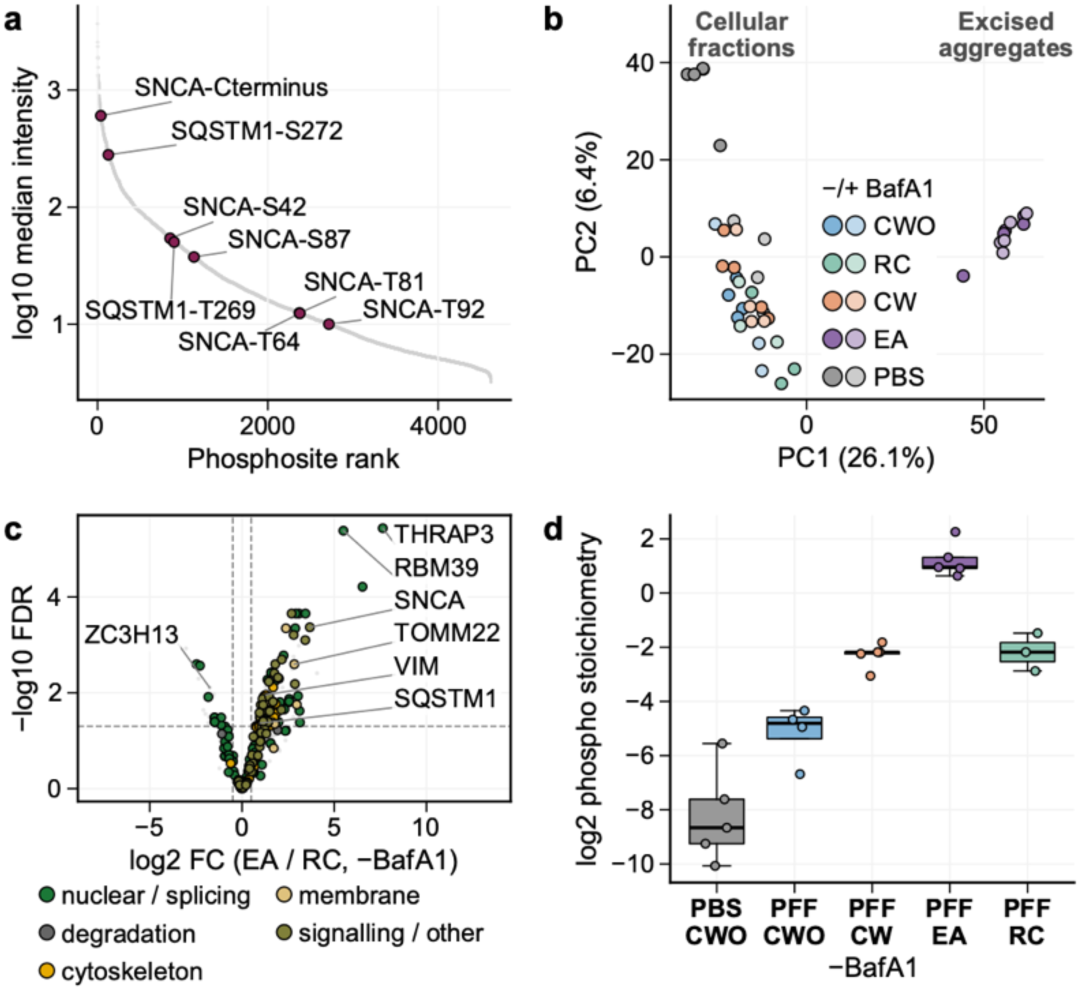
Aggregate DVP phosphoproteomics (nanoPhos) of HEK293 cells expressing A53T α-synuclein. (a) Rank-abundance plot of the 4,622 Class-I phosphosites, ordered by descending median intensity (log scale). α-synuclein (SNCA) and p62/SQSTM1 phosphosites highlighted (gold): SNCA pS42, pS87, pT64, pT81, pT92 and pY125 (labeled SNCA-Cterminus, on the C-terminal tryptic peptide) and p62 pS272 and pT269. (b) PCA of the Class-I phosphosite matrix across all 41 samples that passed quality filtering. Color encodes the fraction (CWO, RC, CW, EA; PBS-treated cells in grey), with two shades per fraction for −BafA1 and +BafA1. Excised aggregates separate along PC1. (c) Volcano plot of phosphosite log2 fold change, excised aggregates versus residual cell body (−BafA1; moderated t-test, limma/eBayes, each site normalized to its parent protein abundance). Dashed lines, |log2FC| > 0.5 and BH-FDR < 0.05; 158 sites increased and 28 decreased. Colored points mark functional groups. (d) SNCA C-terminal phosphopeptide (residues 103–140, any localization; −BafA1): summed intensity of all phosphorylated precursors, proteome-normalized per condition (log2 phosphopeptide minus the per-condition mean SNCA protein level). Box plots across fibril-naive cytosol (PBS CWO, n = 5), PFF cytosol (CWO, n = 4), cells with aggregates (CW, n = 5), excised aggregates (EA, n = 5), and residual cell body (RC, n = 3); points, individual wells.

Selectivity extended to modification and not only to protein identity. Against its matched residual cell body, the aggregate carried 158 increased and 28 decreased phosphosites (five aggregate against three residual cell-body wells; Figure 6c), and 269 against 16 under lysosomal inhibition (Figure S9f), so aggregates are generally enriched in phosphorylation. Fraction-to-fraction comparisons detect only marks present in both, so sites absent from the host fraction are missed. Selecting on detection returned 55 such sites without and 75 with lysosomal inhibition, most of them constitutive or fibril-induced but still detectable outside the aggregate. A third group, 16 and 12 sites, was detected only in the inclusion (Figure S9g, h, Methods), the candidate marks of pathology that a host fraction does not carry.

Among these sites, the α-synuclein C-terminal phosphopeptide was roughly 1,000-fold more abundant in the excised aggregate than in fibril-naive cytosol, with the final 10-fold separating the aggregate from the cell remainder, and the gradient persisted under lysosomal inhibition (Figure 6d, Figure S10a). Diagnostic fragment ions place the increase on Ser129 (Figure S10b-d, Methods). Since the site was normalized to its parent protein, the gradient reflects more phosphorylation and not more α-synuclein.

This 10-fold difference could arise if the enzymes that write the mark were concentrated in the inclusion, but this was not the case. Of 67 α-synuclein-linked kinases and phosphatases, 36 were testable in the aggregate and its residual cell body without lysosomal inhibition (39 in either arm); only 5 differed significantly and none by more than 4-fold, against 10-fold for the α-synuclein site itself (Figure S10e, f; PLK2 was not detected), and about one third of the increase over fibril-naive cells was already present in inclusion-free cells. The data are consistent with α-synuclein carrying the mark before a visible inclusion forms and then being retained. They do not fix when the mark was made, and enzymes present at host level could still act locally. This agrees with reports that fibrils are already pS129-modified before LB-like inclusions form in neurons^21^.

## Discussion

Measured against the cell that formed them, α-synuclein inclusions proved to be organized compartments, selectively concentrating cellular components and differing with the location in which they form. Across a cell line, iDA neurons and patient brain, the inclusion proteome was independent of abundance, dominated by membrane, and taken up as whole complexes and organelle pieces. The α-synuclein it carried was largely phosphorylated before the inclusion became visible. Inclusions therefore capture material that enters with its internal proportions preserved and already modified. This composition bears on how Lewy inclusions have been read, as protective deposits, toxic species or failed-degradation products, each predicting a different content. Of these it supports sequestration and failed degradation together, with uncompensated host depletion on one side and receptor-bound cargo still present on the other.

The markers that first identified these inclusions, ubiquitin, p62/SQSTM1, intermediate filaments and phosphorylated α-synuclein, all reappear in our data, now as concentrations measured against the host cell instead of presences on a section. We detected nearly all canonical constituents reported by earlier studies^21–24,26,27^, but only a quarter were enriched over the matched host neuron, and five enriched proteins appear in no earlier dataset^53^. Because each inclusion shares fixation, post-mortem interval and brain bank with its host neuron, and no protein differed between aggregate-free neurons from patients and controls, neither processing nor disease state is likely to account for the signature.

In both neuronal systems the enriched subset was dominated by presynaptic membrane, and in every donor the round inclusion had a higher ratio of membrane to soluble protein than its host, a proteomic measure of the membrane and organelle crowding seen by electron microscopy in patient tissue^13^ and in fibril-seeded neurons^21^. The enriched set is the presynaptic apparatus itself, so an inclusion in a dopaminergic terminal withdraws the machinery that stores and releases transmitter, where these neurons are most vulnerable. The only machine enriched in all three systems was the V-ATPase, complete with the factors that assemble it. It sits in vesicle and organelle membranes, and α-synuclein binds and clusters synaptic vesicles^57,58^, so an inclusion built from vesicle membrane would carry this composition without concentrating α-synuclein itself, which was only modestly enriched over its host. ER and Golgi proteins were enriched only in HEK293 cells, which stably expressed α-synuclein.

Neuritic and somatic inclusions differed, the neuritic concentrating ubiquitin-dependent quality control and the somatic cytoskeletal and membrane proteins. Since neuritic pathology appears first in this model, the two may mark positions along one path as much as two kinds of inclusion. They may also reflect different compartmental responses to aggregation, quality control in the neurite and an aggresome-like cage in the soma. Aggrephagy receptors also accumulated more under lysosomal inhibition in fibril-exposed cells without an inclusion than in the host cell body of cells with one, so either these cells retained enough aggrephagy capacity to clear material before an inclusion formed, or the inclusion slowed receptor turnover once it had formed. In all three systems, selection acted on assemblies more than on single proteins. Between assemblies, hydrophobicity and low solubility predicted retention, extending earlier work on individual proteins^59,60^ to complexes and organelles. Assemblies that deviate from this rule, such as prefoldin, which delivers non-native proteins to TRiC^61^, are candidates for proteins that act on the inclusion, and those that follow it are candidate passengers.

Because complexes and organelle modules move together, a cell loses groups of related proteins to an inclusion, and pairing each inclusion with its host shows which losses go uncompensated. In HEK293 cells, 40 proteins, mostly Golgi and glycosylation machinery, were depleted from the host and not replaced. Their total level in CW was unchanged or lower than in CWO, so these proteins were captured and some presumably degraded collaterally, and part of their activity is likely to have been lost with them. Only 8 proteins were enriched in the inclusion and raised in the host at the same time, among them SQLE, IFRD1 and ESPL1, short-lived proteins whose levels are set by regulated destruction. The two groups therefore separate material the inclusion captures incidentally from cargo routed for degradation that the inclusion still holds. Sequestration has been linked to toxicity for huntingtin inclusions^60^, and ESCRT-III, enriched here in two systems, is a candidate, since its sequestration by α-synuclein aggregates in this cell line depletes the available pool and potentially compromises endolysosomal repair^62^.

In HEK293 cells, α-synuclein also carried much of its defining modification before it entered the visible inclusion. One third of the rise in the C-terminal phosphopeptide was present in aggregate-free cells, and none of the α-synuclein-linked kinases and phosphatases approached its 10-fold difference across the inclusion boundary, so the mark accumulates where its machinery does not. The modification therefore precedes the visible inclusion, though not necessarily aggregation, since fibril-exposed cytosol can hold sub-visible fibrils and Ser129 phosphorylation has been reported to follow initial deposition^63^. If this holds in neurons and brain, the pS129 stain that defines Lewy pathology marks where modified α-synuclein ends up more than where it is modified. The 16 and 12 sites found only in the aggregate, without and with lysosomal inhibition, are candidates for antibodies that label pathology more sharply.

Our study has limitations. Inclusions were classified on stain, size and shape, which does not resolve the finer morphological classes pathologists distinguish; multiplexed staining would let the same workflow select them. Microdissection also does not separate the inclusion from degradative structures docked on it, so responses to clearance blockade are placed at the inclusion and not necessarily within it. Sixteen donors is a small human cohort, although the paired design balances donor, fixation, post-mortem interval and batch within each comparison. All patients were at neocortical Lewy body stage with comparable Alzheimer co-pathology, and seven of the eight patients were female, so the cohort cannot speak to clinical subtype, disease duration or region of onset.

In conclusion, Aggregate DVP shows α-synuclein inclusions to be selectively organized structures whose composition reflects the location they form in and the clearance state of the cell, consistent with sequestration and failed degradation acting together. Measuring against the host also nominates candidate markers, among them the five enriched proteins absent from earlier datasets and phosphosites found in the aggregate and not in the cell around it, and the paired design shows which functions a cell loses to its inclusion and whether it replaces them. Because Aggregate DVP needs only an inclusion that can be stained, recognized by shape and excised whole, with a comparator from the same sample, it can now be applied across morphologies, regions and disease stages of Lewy pathology, and to tau, TDP-43 and polyglutamine pathology, on tissue that brain banks already hold. The composition of a lesion then becomes a starting point for mechanism and therapeutic targets instead of a description of the end state.

## Methods

### Cell culture

#### HEK293

HEK293T TetOff-SNCA-A53T cells^62^ (RRID: CVCL_D4BW) were cultured in DMEM (Thermo Fisher Scientific, cat. no. 11995073) supplemented with 10% FBS (Thermo Fisher Scientific, cat. no. 1027010), 1% Penicillin/Streptomycin (Thermo Fisher Scientific, cat. no. 15140163), and 2 μg/mL Blasticidin S (Thermo Fisher Scientific, cat. no. A1113903).

Before plating, metal frame slides (PPS, 4 µm; MicroDissect GmbH) were placed cavity down in 10 cm dishes and sterilized under UV light for approximately 1 hour. Slides were then transferred to 4-well plates using sterile tweezers. Cells were then trypsinized with TrypLE Express (Thermo Fisher Scientific, cat. no. 12605036), passed through a cell strainer (Corning, cat. no. 352235), and counted on a Countess 3 cell counter (Thermo Fisher Scientific, cat. no. AMQAX2000). Cells were resuspended in media at a density of 2 × 10⁵ cells per 5 mL and plated onto the frame slides, 5 mL per well.

The following day, PBS and OptiMEM (Thermo Fisher Scientific, cat. no. 31985062) were warmed to 37°C and lipofectamine 3000 (Thermo Fisher Scientific, cat. no. L3000008) was allowed to equilibrate to room temperature. α-Synuclein preformed fibrils bearing an A53T mutation (PFFs; 5 mg/mL stock; see dx.doi.org/10.17504/protocols.io.btynnpve for purification protocol) were thawed and diluted 1:20 in PBS to a final volume of 100–300 μL. Diluted PFFs were sonicated on high for 25 cycles of 5 seconds on and 5 seconds off at 4°C in a BioRuptor Plus (Diagenode, cat. no. B01020001) to break up aggregates into smaller, more nucleation-competent fibrils. A master mix was prepared by combining OptiMEM (600 μL), sonicated PFFs or PBS (240 μL), and lipofectamine 3000 (36 μL). For no-PFF controls, PFFs were substituted with an equal volume of PBS. The master mix was gently vortexed and incubated for 10 minutes at room temperature. The mixture was then added dropwise to cells, with gentle vortexing before each well. The procedure is described in detail at: https://doi.org/10.17504/protocols.io.eq2lyjbbplx9/v1.

48 hours post-treatment, the medium was replaced with 5 mL of medium containing 100 nM Bafilomycin A1 (Invivogen, cat. no. tlrl-baf1) or media containing an equal concentration of DMSO. After 6 hours of BafA1 treatment, cells were washed once with PBS (5 mL), then fixed with 5 mL 4% paraformaldehyde (PFA; Thermo Fisher Scientific, cat. no. 28908) for 10 minutes at room temperature. PFA was removed and replaced with PBS prior to downstream processing.

#### Human dopaminergic iPSCs

##### Expression, purification and fibrillization of human α-synuclein

Recombinant wild-type human α-synuclein was expressed in Escherichia coli using a pT7-7 vector and purified by sequential anion-exchange chromatography and reverse-phase HPLC as previously described^64^. Purified monomeric α-synuclein was diluted in PBS (pH 7.5) to 170–200 µM, filtered through a 100-kDa molecular weight cutoff membrane to remove aggregates, and incubated at 37 °C under constant orbital agitation (1,000 rpm) for 5 days to induce fibrillization. Fibril formation was confirmed by thioflavin T (ThT) fluorescence. For neuronal seeding experiments, fibrils were fragmented by probe sonication on ice (20 s total, 20% amplitude, 1 s on/1 s off) to generate pre-formed fibrils (PFFs)^21^. PFFs were characterized by Coomassie staining and transmission electron microscopy, aliquoted, snap-frozen in liquid nitrogen, and stored at −80 °C until use.

##### Differentiation of NGN2-iPSCs into dopaminergic neuronal cultures (iDA)

Functional iDA neuronal cultures were generated from NGN2-inducible human iPSCs following an optimized protocol described previously^42,65^. NGN2-expressing iPSCs (AIW002-02; Early Drug Discovery Unit (EDDU), McGill University, Canada) were initially converted into induced neurons (iNeurons). Cells were seeded one day before induction (day −1) in mTeSR Plus medium (STEMCELL Technologies, Switzerland) supplemented with the ROCK inhibitor Y27632 (Tocris, Switzerland) to improve cell survival. Neuronal differentiation was initiated on day in vitro 0 (DIV0) by replacing the culture medium with induction medium composed of DMEM/F12 (Life Technologies, Switzerland) supplemented with N2, B27, non-essential amino acids (NEAA), brain-derived neurotrophic factor (BDNF, 10 ng mL⁻¹; Peprotech, United Kingdom), glial cell line–derived neurotrophic factor (GDNF, 10 ng mL⁻¹; Peprotech, United Kingdom), mouse laminin (Life Technologies, Switzerland), and doxycycline (2 µg mL⁻¹; Sigma-Aldrich, Switzerland) to trigger NGN2 expression. The same medium formulation was renewed on DIV1. At DIV2, cells were detached using Accutase and 500,000 cells were replated onto poly-L-ornithine and laminin-coated culture surfaces of PPS slide (4 µm; MicroDissect GmbH). Cells were maintained in Neurobasal medium supplemented with N2, B27, GlutaMax, NEAA, BDNF (10 ng mL⁻¹), GDNF (10 ng mL⁻¹), mouse laminin, and doxycycline (2 µg mL⁻¹). Beginning on DIV3, cultures were exposed to midbrain dopaminergic differentiation medium provided in the STEMdiff Midbrain Neuron Differentiation Kit (STEMCELL Technologies, Switzerland) supplemented with doxycycline (2 µg mL⁻¹) and Sonic Hedgehog (200 ng mL⁻¹) to promote dopaminergic lineage specification. Fresh differentiation medium containing the same supplements was applied on DIV6. At DIV9, the STEMdiff Midbrain Neuron Differentiation media was replaced by the maturation medium from the STEMdiff Midbrain Neuron Maturation Kit (STEMCELL Technologies, Switzerland). From this stage onward, half of the culture medium was replaced weekly with fresh maturation medium, and cultures were maintained under these conditions until experimental use. At this stage, cultures were treated either with human α-synuclein PFF at a final concentration of 500 nM or with an equivalent volume of PBS used as vehicle control, corresponding to the volume of PFF solution added to the culture medium for 56 days (DIV56).

##### Human brain tissue

Post-mortem brain tissue was obtained from two brain banks, the Neurobiobank München (NBM; Munich, Germany) and the Brain and Body Donation Program (BBDP) of the Banner Sun Health Research Institute (Sun City, AZ, USA), and was analyzed as two cohorts that were dissected and acquired as separate batches.

The discovery cohort comprised ten donors. BBDP provided middle temporal gyrus from two donors with PD and two neurologically unremarkable controls. NBM provided frontobasal (orbitofrontal) cortex with the adjoining nucleus accumbens from three donors with Lewy body disease and three neurologically normal controls.

The replication cohort comprised six further BBDP donors, three with PD and three neurologically unremarkable controls, sampled from anterior cingulate cortex.

All tissue was formalin-fixed and paraffin-embedded, and sections were cut at 5 µm. Age at death, sex, post-mortem interval, disease duration, Lewy body stage and density, Braak neurofibrillary stage, Thal phase and the region sampled are listed for each donor in Supplementary Table 1. Ethical approval and consent are stated under Ethics and consent.

### Staining

#### Immunofluorescence staining of HEK293 cells

Cells on membrane slides were permeabilized with 0.1% Triton X-100 in PBS for 5 min, washed briefly with PBS, and blocked with 5% BSA in PBS for 30 min. After a brief PBS wash, slides were incubated with rabbit anti-phospho-α-synuclein (pS129, Abcam, AB51253) primary antibody (1:400) in blocking buffer containing 0.1% Triton X-100 for 1 h at room temperature (400 µL per slide). Slides were washed three times with PBS, then incubated with Alexa Fluor 647-conjugated anti-rabbit secondary antibody (1:400, Thermo Fisher Scientific, A21245) and Phalloidin-Alexa Fluor 488 (1:200, Thermo Fisher Scientific, A12379) in blocking buffer with 0.1% Triton X-100 for 1 h at room temperature. After two PBS washes, nuclei were counterstained with Hoechst 33342 (1:1,000 in Milli-Q water, 5 min). Slides were washed twice briefly with PBS and dried by centrifugation at 100 × g for 5 min.

Ferritin immunostaining was performed on HEK293T cells grown on poly-L-lysine-coated coverslips (Neuvitro, GG-12-1.5-PLL). Cells were fixed in 4% PFA (Thermo Fisher Scientific, 28908) for 10 min at room temperature and subsequently permeabilized with 0.1% Triton X-100 in PBS for 5 min at room temperature. A 1 hr blocking step followed, using a Blocking Solution of PBS containing 5% milk powder (Sucofin) and 0.1% Triton X-100 at pH 7.4. Primary antibodies, diluted in this Blocking Solution, were applied to the coverslips and left overnight at 4°C at the following concentrations: mouse anti-pS129 α-syn at 1:1000 (Abcam, ab184674) and rabbit anti-Ferritin light chain at 1:1000 (Proteintech, 10727-1-AP). Following three washes in PBS supplemented with 0.02% Triton X-100, coverslips were incubated with secondary antibodies (1:1000 in Blocking Solution) for 3 hr at room temperature in the dark.

The secondary antibodies used were Alexa Fluor 568 Goat anti-Mouse IgG (H+L) Cross-Adsorbed (Thermo Fisher Scientific, A-11019) and Alexa Fluor 488 Goat anti-Rabbit IgG (H+L) Cross-Adsorbed (Thermo Fisher Scientific, A-11008). The actin cytoskeleton was then labeled with Phalloidin-iFluor 647 Reagent (Abcam, ab176759) at 1:1000 in PBS containing 1% BSA and 0.02% Triton X-100 (pH 7.2) for 45 min. Nuclei were counterstained with NucBlue (Thermo Fisher Scientific, R37606) for 5 min, after which two further PBS washes were performed. Coverslips were finally mounted onto Epredia glass slides (Thermo Fisher Scientific, 17294884) using Dako Fluorescence Mounting Medium (Agilent, S3023). The full protocol is available at https://doi.org/10.17504/protocols.io.e6nvwd7n7lmk/v1.

#### Immunocytochemistry of α-synuclein pathology formed in iDA neurons

iDA neuron cultures grown on PPS membrane slides were rinsed twice with phosphate-buffered saline (PBS) and fixed in 4% formaldehyde (Sigma-Aldrich, Switzerland) for 20 min at room temperature. Cells were then permeabilized and blocked for 30 min at room temperature in PBS containing 0.1% Triton X-100 (PBS-T) and 3% bovine serum albumin (BSA; Sigma-Aldrich, Switzerland). Following this step, cultures were incubated overnight at 4°C with primary antibodies: mouse anti-phospho-α-synuclein pS129 (clone 81A; BioLegend, 825701; RRID: AB_2564891; 1:2,000), chicken anti-β-tubulin III (Merck, AB9354; RRID: AB_570918; 1:2,000), and rabbit anti-p62/SQSTM1 (Proteintech, 18420-1; RRID: AB_10694431; 1:500). After primary antibody incubation, cells were washed five times with PBS-T and incubated with the appropriate fluorescent secondary antibodies: donkey anti-rabbit Alexa Fluor 488 (Invitrogen, A21206; 1:800), donkey anti-mouse Alexa Fluor 568 (Invitrogen, A10037; 1:800) and donkey anti-chicken Alexa Fluor 647 (Jackson ImmunoResearch, 703-605-155; 1:400), together with DAPI (Life Technologies, Switzerland) to label cell nuclei. Samples were washed five additional times with PBS-T, rinsed twice with PBS, and stored in PBS until imaging. As previously optimized, pathological α-synuclein accumulation was assessed using the phosphorylation-specific pS129 antibody, which selectively detects aggregated α-synuclein species^42^. Neuronal morphology and neuritic architecture were visualized by co-immunostaining for β-tubulin III, enabling identification of neuronal somata and processes. Nuclei were counterstained with DAPI.

For the validation of proteins identified by DVP, iDA cultures grown on coverslips and fixed in 4% formaldehyde were stained in a similar manner. Cells were permeabilized and blocked in PBS-T for 30 min, and then incubated overnight at 4°C with the primary antibodies: anti-UBQLN2 (clone EPR28592-74; Abcam, ab323454; 1:500), anti-proSAAS (Sigma Aldrich; ABN2268; 1:100) or anti-Chromogranin A antibody (clone RM1025; Abcam; ab283265; 1:100) in combination with anti-phospho-α-synuclein pS129 (clone 81A; BioLegend, 825701; RRID: AB_2564891; 1:2,000) and chicken anti-β-tubulin III (Merck, AB9354; RRID: AB_570918; 1:2,000). The cells were then washed and incubated for two hours at room temperature with the secondary antibodies donkey anti-mouse Alexa Fluor 568 (Invitrogen, A10037; 1:800), donkey anti-chicken Alexa Fluor 647 (Jackson ImmunoResearch, 703-605-155; 1:400), and donkey anti-rabbit Alexa Fluor 488 (Invitrogen, A21206; 1:800), together with DAPI (Life Technologies, Switzerland). The coverslips were then washed five times in PBS-T before being rinsed in H2O and mounted onto glass slides with polyvinyl alcohol mounting medium containing DABCO (Sigma; 10981).

#### Immunohistochemistry and cresyl violet staining of FFPE brain tissue

To prepare slides for laser microdissection-compatible staining, PEN membrane slides (2 µm; MicroDissect GmbH) were activated under UV light (254 nm, 1 h) and functionalized with Vectabond tissue adhesive (Vector Laboratories; SP-1800-7) according to the manufacturer’s instructions. Sections were mounted onto the coated membrane slides and stored at room temperature until further processing.

Sections were baked at 55°C for 30 min, deparaffinized with xylene (2 × 2 min), and rehydrated through a graded ethanol series (100%, 90%, 75%, 30%, each 2 × 1 min) followed by Milli-Q water (2 × 1 min). Antigen retrieval consisted of 75% formic acid applied directly on the slide for 20 min, followed by Milli-Q water washes (2 × 2 min) and heat-induced antigen retrieval in Dako pH 6 buffer (DAKO, S2369) with 10% glycerol at 88°C for 40 min. After cooling for 5 min and Milli-Q water washes (2 × 2 min), endogenous peroxidase activity was quenched with 3% hydrogen peroxide in water for 20 min at room temperature in the dark. Slides were washed in Milli-Q water (1 × 2 min) and PBS (2 × 2 min), then blocked with 0.1% BSA in PBS for 60 min at room temperature. Sections were incubated overnight at 4°C with rabbit anti-phospho-α-synuclein (pS129, Abcam, AB51253) primary antibody (1:1,000) in antibody diluent with 0.1% BSA in a wet staining chamber. After PBS washes (2 × 2 min), sections were incubated with goat anti-rabbit IgG HRP (1:400, Thermo Fisher Scientific, G21234) in PBS with 0.1% BSA for 60 min at room temperature, washed in PBS (3 × 2 min), and developed with a 1xDAB substrate working solution (metal enhanced DAB substrate kit, Thermo Scientific, 34065, DAB/Metal concentrate (10x) diluted in Stable Peroxide Substrate Buffer to a 1x working solution) for 2-10 min with microscopic monitoring of signal development. Slides were washed in PBS (2 × 2 min).

For counterstaining, sections were dehydrated in 70% ethanol for 5 min, rinsed in Milli-Q water, and stained with 0.2% cresyl violet acetate (MP Biomedicals, 150727) in Milli-Q water containing 0.4% ethanol for 10 min at 30°C. After rinsing in Milli-Q water (2×), differentiation was controlled by brief immersion (1-2 dips, maximum 15 s) in an ethanol-acetic acid solution (50 mL ethanol, 4 drops acetic acid) with microscopic monitoring. Sections were rinsed in 100% ethanol (1-2 dips), dehydrated in 100% isopropanol (2 × 5 min), and air-dried.

### Microscopy

#### Fluorescence microscopy of HEK293 cells and iDA neurons

Stained membrane slides were imaged without coverslips on a PerkinElmer Opera Phenix high-content screening system controlled by Harmony v5.2 software, using a 20× air objective (NA 0.8) in confocal mode. Images were acquired at 2160 × 2160 pixel resolution with 1× binning, a resulting pixel size of 0.29899 µm/pixel and 10% overlap between adjacent fields of view. A single focal plane was recorded per slide, with the z-position adjusted manually for each channel to account for chromatic focal shifts. Up to four channels (DAPI, Alexa 488, Alexa 568, Alexa 647) were acquired sequentially to prevent spectral crosstalk, with exposure times ranging from 60 to 600 ms and laser power set between 60% and 100% depending on the channel.

Confocal images were collected at the Max Planck Institute of Biochemistry Imaging Facility (RRID:SCR_025739) using a Leica SP8 Falcon equipped with a 63X oil objective and operated through Leica Application Suite X 3.5.7.23225 (Leica). Figure panels were generated with a custom MATLAB script (https://github.com/csitron/Microscopy_lif_Adjust_MATLAB). Intensity adjustments were applied uniformly across each entire image, and identical minimum and maximum cutoff values were used whenever images were compared directly.

#### BrighRield microscopy of human brain tissue

Immunohistochemically stained and cresyl violet-counterstained FFPE sections were coverslipped and digitized on a Zeiss Axioscan 7 slide scanner in brightfield mode using a transmitted light LED lamp. Whole-slide images were acquired with a Plan-Apochromat 20×/0.8 NA M27 objective and an Axiocam 705 color camera at 24-bit RGB, with 1× binning and 10% tile overlap, yielding a pixel size of 0.172 × 0.172 µm. No z-stacks were acquired. Illumination settings were calibrated on one section and applied consistently across all samples within each experimental group. Tiles were stitched using Zeiss ZEN imaging software. After imaging, coverslips were removed by submerging slides face-down in Milli-Q water until the coverslip detached. Slides were subsequently air-dried before laser microdissection.

### Image analyses

#### HEK293 cells and iPSCs

For both cell culture models, tiled images were stitched using scPortrait^66^ and subsequently analyzed in BIAS (Single Cell Technologies; https://www.sct.bio, v2025-03-19). Images were imported via the XXL image import module at a resolution of 0.29899 µm/pixel and retiled with 10% overlap. Tiles lacking cellular content and border regions were excluded prior to downstream analysis.

For HEK293 cells, selected tiles were exported for segmentation using Cellpose 2^44^ with custom-trained models to generate separate masks for cell bodies and aggregates. After re-importing masks into BIAS, size filtering and artifact removal were applied to refine segmentation quality. Cells were classified as aggregate-containing or aggregate-free based on mask overlap, and the resulting cell and aggregate compartment classes were exported as XML files for laser microdissection.

For iDA neurons, selected tiles were exported and processed with a custom CellProfiler (v4.2.8)^67^ pipeline to generate soma masks: β-tubulin images were smoothed with a Gaussian filter (diameter 4 pixels) and processed by morphological opening (disk element, 20 pixels) to separate neurite from somatic signal. The neurite-subtracted image was segmented using global 2-class Otsu thresholding (minimum diameter 40 pixels) to identify neuronal cell bodies. The resulting binary masks were exported and imported into BIAS. Aggregates, p62 signal, and β-tubulin signal were segmented by intensity thresholding within BIAS. Soma and neurites were classified as aggregate-containing or aggregate-free. Aggregates were further filtered by p62 positivity and association with intact cells; only p62-positive aggregates located within cells were retained for further analysis. These were then classified into round (somatic) and neuritic aggregates using the clustering module in BIAS. Final compartment classes were exported as XML files for laser microdissection.

#### Human brain tissue

Images were converted to spatialdata objects^68^ with dvp-io version 0.4.2. Whole-slide images were color-deconvolved into their 3,3ʹ-diaminobenzidine tetrahydrochloride and cresyl violet (CV) channels using the H-DAB color deconvolution matrix^69^ implemented in scikit-image^70^ (skimage.color.separate_stains). Binary masks for each stain were generated by intensity thresholding of the deconvolved channels, with DAB-positive pixels defined as a deconvolved intensity between 0.42 and 0.85, the lower bound set at the 99.9th percentile of the DAB channel. Downstream mask processing was performed as follows. DAB-positive labeled masks were filtered to an area of 10 to 400 µm², removing small artifacts and large confluent staining, and vectorized into polygon geometries. Solidity was computed as the ratio of object area to the area of its convex hull, and eccentricity from the best-fit ellipse. Objects with solidity ≥ 0.6 and eccentricity ≤ 0.9 were classified as round and all others as elongated. Cresyl violet–positive masks were similarly area-filtered, vectorized, and classified as likely glial cells or likely neurons by computing a composite neuron score derived from the area, circularity, eccentricity, and solidity of each object. Neuropil regions were circles placed at random in areas free of DAB-positive and cresyl violet-positive masks. The obtained contours were exported to XML format using dvp-io for downstream spatial sampling.

### Laser microdissection

The amount of material collected per sample was defined by excised area. For each system a reference object area was set, and the number of shapes collected per sample was chosen so that the total excised area equaled a fixed number of reference-sized objects. In HEK293 cells the reference was 600 µm², the mean area of a whole cell in this dataset, and 100 reference-sized objects were collected per sample and 350 for nanophos experiments. The same 600 µm² reference, defined in the HEK293 dataset, was applied to iDA neurons, where 50 reference-sized objects were collected per sample. In brain tissue the reference was 90 µm², the mean area of a neuron in that dataset, and 100 reference-sized objects were collected per sample. Because the dissected classes differ in size, a class of smaller objects contributed more shapes per sample than a class of larger ones, for example excised aggregates against whole cells, while the excised area per sample was held constant.

XML-formatted contour outlines were registered to reference points and loaded into a Leica LMD7 system running LMD software v8.5. All cutting was performed with 63× and 40× objectives in semi-automated mode (aperture 1, speed 100, middle pulse count 1, final pulse 1, pulse frequency 2,900). Offset was set to 190 for cell culture and 100 for brain tissue; head current and power were adjusted between 45 and 60% per sample. Under these settings, cutting throughput reached approximately 1,000 shapes per hour. Excised shapes are collected by gravity into 384-well collection plates (Eppendorf; 0030129547), with collection restricted to alternating wells excluding the outermost rows and columns to avoid cross-contamination. A UV shield mounted above the sample stage minimized airflow-induced sorting errors. After collection, plates were sealed, spun at 1,000 × g for 10 min, and stored at −20°C.

Laser microdissection has a nominal optical resolution of ∼200 nm; on fixed tissue the effective precision of an excised contour is closer to 1 µm. The compact inclusions profiled here, 50–100 µm² in cross-section (≈8–11 µm across), exceed this precision by roughly an order of magnitude and were excised as discrete objects, whereas the thin neuritic (∼20 µm²) and the still smaller elongated brain inclusions approach this floor in their narrow dimension.

### Sample preparation for proteomics samples

Lysis and digestion were carried out following an automated protocol adapted from Rosenberger *et al.*^38^. Collection plates were thawed and 28 µL of acetonitrile (ACN) was dispensed into each well using a Bravo pipetting robot (Agilent) to recover excised material from the well walls. Plates were then vacuum-dried at 45°C in a SpeedVac. All subsequent dispensing steps were performed on a Mantis liquid handler (Formulatrix). Dried samples were resuspended in 7 µL of 70 mM triethylammonium bicarbonate (TEAB, pH 8.5) containing 0.013% n-dodecyl-beta-D-maltoside and heated at 95°C for 1 h. Following addition of 1 µL ACN, a second incubation at 75°C for 1 h was performed. Enzymatic digestion was initiated by adding 2 µL of a combined trypsin/Lys-C solution (6 and 4 ng/µL, respectively, in 70 mM TEAB) and allowed to proceed overnight (16 h, 37°C). Reactions were stopped with trifluoroacetic acid (1% final concentration).

### Sample preparation of phosphoproteomics samples

#### Sample lysis

Collection plates containing the various numbers of shapes were thawed and 28 µL of ACN was added to each sample-containing well using Bravo pipetting robot, after which plates were spun down at 3000 g for 10 min to recover excised material from the well walls. Plates were then dried at 30 °C in a SpeedVac. 10 µL of a lysis buffer (2% sodium deoxycholate, 0.1% DDM, 10 mM tris(2-carboxyethyl)phosphine, 40 mM chloroacetamide in 100 mM Tris-HCl pH 8.5) was added to each well and the plate was then centrifuged for 1 min at 3000 g. Sample lysis was then performed in a 3-step manner: samples were first heated at 95°C for 15 min, followed by a high-energy sonication step for 10 min (10 sec on/off pulses) after which a second heating step was performed at 95°C for 15 min. Sample lysate was then split in the proportion of 1:9 into proteome:phosphoproteome sample preparation branches.

#### SPEC and nanoPhos sample preparation

Proteome and phosphoproteome samples were prepared using SPEC and nanoPhos sample preparation workflows, respectively^35,56^. Briefly, SPEC tips were prepared in-house by placing two layers of strong anion exchange material (3M Empore) in a pipette tip with a blunt-end syringe needle. Tips were activated using 10 µL of dimethyl sulfoxide and centrifuged at 700g for 1.5 min. Next, tips were preequilibrated with 20 µL of SPE preequilibration buffer (20 mM 3-(Cyclohexylamino)-1-propanesulfonic acid (CAPS) pH 10.5, 0.01% DDM in 50% methanol) by centrifugation at 700g for 2 min. Then, 1 µL (proteome) or 9 µL (phosphoproteome) of sample lysate was loaded onto SPEC tips in SPEC equilibration buffer at a 1:45 (proteome) or 1:10 (phosphoproteome) lysate:buffer ratio and centrifuged at 400g for 10 min. The tips were then washed using 20 µL SPEC washing buffer (50 mM TEAB, 0.01% DDM) by centrifuging at 700g for 3 min or until all liquid has passed through the tip. Protein digestion was performed by applying 2 µL of digestion buffer (25 ng/ µl trypsin, 25 ng/ µl LysC, 50 mM TEAB, 0.01% DDM) into the tip, centrifuging at 100g for 20 sec and incubating at 37 °C for 90 min. Proteome samples were eluted from SPEC tips by applying 20 µl of SPEC elution buffer (1% formic acid, 0.01% DDM) directly into preequilibrated Evotip Pure C18 tips by centrifuging at 300g for 5 min. For phosphoproteome analysis peptides were eluted by the addition of 20 µl of nanoPhos Elution buffer (1M NaCl, 10% ACN, 0.1% DDM, 1% FA in ddH2O) to a 96-well TwinTec Eppendorf plate, which already contained 80 µl of nanoPhos preelution buffer (90% ACN, 0.1% DDM, 1% FA in ddH2O).

#### Phosphopeptide enrichment for nanoPhos experiments

Phosphopeptide enrichment cartridges, each containing 5 µl Fe(III)-nitrilotriacetic acid (Agilent, part number: G5496-60085), were first primed with 100 µl priming buffer (1% FA, 99% ACN) with a flow rate of 300 µl/min, followed by equilibration with 50 µl wash/equilibration buffer (1% FA, 80% ACN in ddH_2_O) with a flow rate of 5 µl/min. Peptides were then loaded on cartridges with a flow rate of 5 µl/min and subsequently washed with 50 µl wash/equilibration buffer with a flow rate of 5 µl/min. Phosphopeptides were eluted with 25 µl of elution buffer (500 mM NH_4_H_2_PO_4_ in ddH_2_O) with a flow rate of 5 µl/min directly onto preequilibrated EvoTips.

### Evotip loading

Digested peptides were loaded directly onto Evotips without intermediate freezing. Tips were activated by soaking in 1-propanol (5 min), washed twice with 50 µL buffer B (ACN, 0.1% formic acid), soaked a second time in 1-propanol (5 min), and conditioned with a single 50 µL buffer A wash (0.1% formic acid). Peptide samples were then applied in 70 µL buffer A pre-loaded on the tip. The C18 disk was topped with 150 µL buffer A and briefly centrifuged (700 × g, 1 min) to seat the peptides. Tips were kept in buffer A until measurement.

### LC-MS/MS acquisition (proteomics)

Peptide samples were analyzed on an Evosep Eno liquid chromatography system (Evosep) coupled online to an Orbitrap Astral mass spectrometer (Thermo Fisher Scientific). Separation was performed using the Evosep low-flow Whisper Zoom gradient at a throughput of 80 samples per day on an Aurora Rapid 5 × 75 C18 UHPLC column (IonOpticks), maintained at 50°C with a column heater (IonOpticks). Ionization was achieved through a FAIMS Pro interface and an EASY-Spray source (both Thermo Fisher Scientific) at an electrospray voltage of 1,900 V and a FAIMS compensation voltage of −40 V with a total carrier gas flow of 3.5 L/min. FAIMS electrode temperatures were set to 100°C (inner) and 80°C (outer).

Full MS1 survey scans were recorded in the Orbitrap from 380 to 1,500 m/z at 240,000 resolution (at m/z 200) with a normalized AGC target of 500% and a maximum injection time of 100 ms (RF lens 40%). Data-independent acquisition on the Astral analyzer employed 75 variable-width isolation windows spanning 380-1480 m/z, designed using py_diAID^71^ with fifteen further windows of 2 Th centered at fixed m/z values between 964 and 1494 (Supplementary Table 2) to match precursor density across the mass range. Fragment ions were generated by higher-energy collisional dissociation at 25% normalized collision energy and scanned from 150 to 2,000 m/z with an AGC target of 500% and a maximum injection time of 10 ms.

### LC-MS/MS acquisition (phosphoproteomics)

Phosphoproteomics samples were analyzed using an Evosep Eno LC system (Evosep) coupled to an Orbitrap Astral Zoom mass spectrometer (Thermo Fisher Scientific). Peptides were eluted from the Evotips using a Whisper Zoom gradient with a throughput of 80 samples per day on an Aurora Rapid column of 5-cm length, 75-µm-internal diameter, packed with 1.7 µm C18 beads (IonOpticks). The column temperature was maintained at 60 °C using a column heater (IonOpticks). The Orbitrap Astral Zoom was equipped with an EASY-Spray source (Thermo Fisher Scientific). An electrospray voltage of 1,900 V was applied for ionization, and the radio frequency level was set to 40. For phosphoproteomics samples Orbitrap MS1 spectra were acquired from 380 to 1,580 m/z at a resolution of 240,000 (at m/z 200) with a normalized automated gain control (AGC) target at 500% and a maximum injection time of 50 ms. For the Astral MS/MS scans in data-independent acquisition (DIA) mode, we used 100 variable isolation windows, designed with a pyDIAid software^71^ (Supplementary Table 3). A maximum injection time of 10 ms was used. The isolated ions were fragmented using high-energy collisional dissociation with 25% normalized collision energy. The proteome fraction prepared in parallel from the same samples (the 1:9 proteome branch of the split lysate) was acquired on the same instrument: Orbitrap MS1 spectra from 380 to 980 m/z at a resolution of 240,000 (at m/z 200), a normalized AGC target of 500%, and a maximum injection time of 3 ms; for the Astral MS/MS scans in DIA mode, 200 equidistant isolation windows of 3 Th each were used with a maximum injection time of 6 ms, and isolated ions were fragmented by higher-energy collisional dissociation at 25% normalized collision energy. Samples were acquired in randomized order within each condition.

### Spectral search

Spectral libraries were built in two steps. A library predicted in silico from a canonical human FASTA database (20,435 entries, UniProt, downloaded 4 April 2024), with trypsin/P specificity and up to one missed cleavage, was used for a first DIA-NN search, which produced a project-specific empirical spectral library. For HEK293 cell searches, the A53T α-synuclein mutation was manually introduced into the FASTA sequence. Raw files were searched with DIA-NN v2.3.2^72,73^ against a project-specific empirical spectral library, with a human canonical FASTA (20,416 entries) supplied for protein annotation. Trypsin/P specificity was used with up to one missed cleavage, and N-terminal methionine excision was enabled. Methionine oxidation (UniMod:35) and protein N-terminal acetylation (UniMod:1) were set as variable modifications, with a maximum of three variable modifications per peptide. Peptide length was restricted to 6-40 amino acids, precursor charge states to 1-5, precursor m/z to 300-1,500, and fragment ion m/z to 200-1,800. MS1 and MS2 mass accuracies were fixed at 5 and 10 ppm respectively, with a scan window radius of 7. Peptidoform-level scoring and IDs, RT and IM profiling refinement were enabled, relaxed protein inference was disabled, and library precursors were not reannotated against the FASTA. Searches were performed in two passes: each file was first searched individually, after which all files of a dataset were reanalysed together with match-between-runs, reusing the first-pass quantities and generating a combined spectral library. Precursors were filtered at 1% FDR. All remaining parameters were left at their DIA-NN 2.3.2 defaults.

Phosphoproteomics LC-MS raw files were processed in Spectronaut v21.1 (build 21.1.260825.94842) without experimental spectrum libraries (directDIA+ (Deep) workflow). A canonical human FASTA database (20,416 entries, UniProt, downloaded 22 May 2026) with the manually introduced A53T α-synuclein mutation was used for spectral library prediction. We set the protease specificity to trypsin (Trypsin/P, specific digestion) with a maximum number of two missed cleavages and a peptide length of 7-52 amino acids. The mass tolerances for precursor and fragment ions were set to ‘Dynamic’ for both MS1 and MS2 level. False discovery rates were controlled by a target-decoy approach to <1% at precursor and protein levels. For phosphoproteomics experiments, we defined cysteine carbamidomethylation as a fixed modification and protein N-terminal acetylation, methionine oxidation and serine/threonine/tyrosine (STY) phosphorylation as variable modifications (maximum five variable modifications per peptide) in ‘BGS Phospho PTM Workflow’ with PTM localization cutoff set to 0. Cross-run normalization option was enabled and set to ‘Automatic’. For the proteomics part of the phosphoproteomics experiment we used ‘BGS Factory Settings (default)’ workflow with default settings.

### Data analysis

Analyses used Python 3.12 and alphapepttools 0.2.0 (https://github.com/MannLabs/alphapepttools)^74^. The notebooks, the contaminant, excluded gene set and curated gene lists, the gene set decisions and the rules of their AI review are published with the code at https://github.com/MarcMarc94/aggregateDVP_analysis. P-values were adjusted by the Benjamini-Hochberg (BH) procedure within each contrast, except in gene set enrichment, where the false discovery rate (FDR) is the permutation q-value of gseapy. Significance means FDR < 0.05, and fold changes are log2 (log2FC). A replicate is one laser-microdissected pool from one culture in the cell models, and the donor is the unit of analysis in brain tissue.

### General procedures

#### Preprocessing and testing

Protein-group intensities from the DIA-NN protein group matrix were log2(*x* + 1) transformed without further normalisation. Samples with fewer than 2,000 protein groups were excluded. Keratins, protein groups with only cRAP identifiers, immunoglobulins (gene symbols starting with IGK, IGL or IGH, except the IgLON family) and sample-type-specific serum, skin and laboratory proteins were removed. A protein group’s gene symbol is the first of its DIA-NN annotation. Values were not imputed unless stated. The coefficient of variation is the standard deviation (*n* − 1 denominator) over the mean of linear intensities, for proteins with at least three values per condition (in brain tissue, per bank and cell type). Principal component analysis (PCA) used mean-centred, unscaled log2 intensities of the protein groups quantified in every sample. The cell models were tested by Welch’s t-test (at least two values per group), and brain tissue and phosphosites by limma with empirical Bayes moderation (inmoose) unless stated. For compartment composition, each compartment is a union of selected GO cellular component sets (MSigDB, retrieved 8 September 2026), one of them named Other, and each protein was assigned to the smallest compartment containing it or else to Other. Linear intensities, from condition means of log2 values in the cell models and per pool in brain tissue, were summed per compartment as a percentage of the total, and brain percentages were averaged per donor and fraction.

#### Gene set enrichment

Gene set enrichment analysis (GSEA) used gseapy prerank (1,000 gene permutations, seed 42) on MSigDB sets with 20 to 500 ranked genes, obtained through decoupler. The region-paired brain fit was tested against the same GO cellular component sets, all other analyses against Reactome and GO biological process (retrieved 31 July 2026). Genes were ranked by Welch’s *t* from measured values (at least three per group), and a symbol shared by several protein groups was represented by the group with the fewest missing values, then the highest abundance. Three rankings differ and use the most abundant protein group per symbol. The HEK293 BafA1-induced shift (Figure S3c) was ranked by its value, the cross-system brain contrast by the moderated *t* and the region-paired brain fit by the signed normal quantile of the moderated-*t* p-value. Sets with a normalized enrichment score (NES) and p-value of exactly 1, which occur when every permutation reproduces the observed score, were removed. Sets naming a cell type or tissue absent from the system, and generic containers such as cytoplasm in brain tissue, were tested and kept in the full result tables but removed before clustering. Significant sets were clustered within each direction by the overlap coefficient of their leading edges (shared genes over the smaller leading edge), with complete linkage so that every pair in a cluster has a coefficient of at least 0.5. For the 40 clusters per direction with the largest absolute NES (20 in brain tissue), the authors chose a representative set and a label from the genes shared by the leading edges, assisted by the AI review (Claude, Anthropic). In multi-contrast figures each row is one label and shows one gene set in every column, the label’s representative with the largest absolute NES (HEK293, cross-system) or the set chosen by the gene-list rule (iDA neurons). Rows were admitted when significant in at least one contrast, placed in the direction of their strongest significant cell and filled by the largest absolute NES, after three rows per contrast and direction in iDA neurons.

### HEK293

#### Fractions and protein selections

Protein groups quantified in all six replicates of at least one condition were tested (7,013). The PFF-against-PBS volcano (Figure 2b) marks FDR < 0.05 and |log2FC| > 0.3. The four PFF fractions were compared by one-way ANOVA (scipy, at least two values per group), with and without BafA1 separately, and the ANOVA-significant proteins by two planned Welch’s t-tests with BH correction among those proteins. The tests compared rest cells (RC) with cells without aggregates (CWO) and excised aggregates (EA) with cells with aggregates (CW). Each test called a protein higher, lower or unchanged (FDR < 0.05, |log2FC| > 0.3), giving nine redistribution categories. The aggregate composition dot plot (Figure 2f) shows the 15 proteins with the largest positive log2FC (FDR < 0.05) in each of EA against RC with and without BafA1 and the BafA1 effect in EA, plus the proteins in at least two of the corresponding top-50 lists, 47 proteins in all, grouped by function by hand. BafA1-responsive proteins are significant in at least two of the five BafA1 contrasts, have one sign in every contrast with a value and a largest |log2FC| of at least 0.65 (49 proteins). Figure 2g shows 46 of them (IREB2, NCOA4 and FTL are in Figure S3e) with SQSTM1, TAX1BP1, MAP1LC3B and the controls ACTB, GAPDH, TUBB, H4C1 and PSMA1.

#### Excision purity and mass balance

Purity places each sample on a scale from CWO (0) to EA (100), built from the 24 wells of the −BafA1 arm. The linear intensities of the 5,730 proteins quantified in all 24 wells were divided by each well’s sum. Markers differed between EA and CWO by Welch’s t-test on log2 compositions (FDR < 0.05, |log2FC| ≥ 1) and twofold between the mean compositions of the two anchors (1,124 markers). A well’s position is the median over markers of (*x* − CWO)/(EA − CWO), with *x* its composition. CW and RC wells were placed on the scale built from all anchor wells, CWO and EA wells by leave-one-out from the other eleven anchor wells, and the means of the leave-one-out placements define floor (CWO) and ceiling (EA). For mass balance, cells with aggregates were modelled as CW = *w* × EA + (1 − *w*) × RC. Seven misspecified models replaced the host by CWO or PBS-treated cells or the target by RC or CWO, removed one component with *w* fixed, or permuted CW across proteins. All models used the 5,700 proteins quantified in the 30 −BafA1 wells of the five fractions and their 1,113 markers. For components *A* and *B*, *w* is the median over markers of (target − *B*)/(*A* − *B*). The residual, the median absolute log2 ratio of observed to predicted composition, was divided by 0.6745 times the median over markers of its per-protein standard deviation in 200 bootstrap resamplings of the six wells per fraction.

### iDA neurons

#### Matrices, tests and selections

Coefficients of variation used the protein groups quantified in all replicates of at least one condition, and PCA, t-tests and GSEA those quantified in every sample after contaminant removal. Only for Figure S4c, proteins quantified in at least 80% of the replicates of one condition were kept and their missing values drawn from a Gaussian distribution with a mean 1.8 standard deviations below the protein’s mean and a width of 0.3 standard deviations, both over all samples (seed 42). The markers of Figures 3c and S4c are curated nuclear and process markers, and the volcano and the marker match use FDR < 0.05 and |log2FC| > 0.5. The ubiquitin-dependent quality control panel (Figure 3g) used all protein groups after contaminant removal. A protein entered a contrast when both groups had values in at least 50%, and at least two, of their samples, with BH correction over all proteins tested in that contrast. For Figure S4d, a gene list was written by hand for each label from its name before any leading edge was read. Each tested set was scored by purity (fraction of its leading edge in the list) and coverage (fraction of the list in its leading edge). Candidates have purity ≥ 0.45 and coverage ≥ 0.10 and qualify when their purity reaches 0.30 in every contrast in which they are significant. The qualifying set with the highest purity in a significant contrast, then the largest |NES| in either contrast, fills both columns. The PFF response heatmap (Figure 3d) shows hand-chosen leading-edge proteins of the sets in Figure S4d and five PD-linked proteins, and dense-core vesicle proteins come from a curated list. The aggregate composition heatmap (Figure 3f) shows the 15 most enriched and 5 most depleted proteins (FDR < 0.05) of each of its three contrasts, grouped by function.

### Post-mortem human brain

#### Inclusions, hosts and donors

Of 385 microdissected pools, four were excluded for cutting problems, 12 were unassigned random controls and 37 of the remaining 369 had fewer than 2,000 protein groups. Contaminant removal left 6,842 of 6,941 protein groups, and those quantified in at least 80% of the pools of one fraction were kept (4,643). Round inclusions were compared with their DAB-positive host neurons in a region-paired design. Within each donor and region, the mean log2 intensity of the host pools was subtracted from that of the inclusion pools, with the cortical samples of both banks as one region, and the region differences were averaged per donor without weights (seven donor-region pairs, 2 to 5 inclusion pools each). The 3,660 proteins with a value in every donor were fitted by an intercept-only limma model (prior 3.0 and residual 4 degrees of freedom). Aggregate-free (DAB-negative) neurons of disease and control donors were compared by limma on donor means over all regions, for proteins with a value in every donor. The membrane-to-nuclear ratio of a pool is the log2 ratio of the summed linear intensities of two fixed GO cellular component groups, membrane and vesicle against soluble nuclear and translation, without genes in both, and was averaged per donor and fraction. In Figure 4f a donor’s value is its mean difference over the detected members of the representative set. Direction consistency counts the significant sets, without wrong-lineage sets and generic containers, in which every donor’s value minus its mean over all other proteins has the sign of the NES.

#### Variance between patients and replication

Variance was partitioned in the only stratum with replicate inclusion pools in several donors, the nucleus accumbens of the Munich bank (three donors, 13 inclusion pools, 12 host wells, 1,688 proteins quantified in all). Each pool was expressed as its log2 ratio to its own donor’s mean DAB-positive neuron in that region. The total sum of squares about the grand mean was split into the protein main effect and a remainder, and the remainder into between-patient, within-patient and donor-mean sampling components, estimated per protein by the method of moments (observed mean squares set equal to their expected values). Negative between-patient estimates were summed as obtained, and the between-patient component is reported as a share of the remainder. Per-protein patient differences were tested by exact permutation, comparing the range of the three donor means with its values under all distinct assignments of the 13 pools to the three donors with the observed group sizes, also after regressing out the number of protein identifications per pool. For replication, both cohorts were searched together in DIA-NN (534 pools before filtering). Inclusion-over-host differences were computed as above, without merging cortical regions, for five discovery and three replication donors, on protein groups quantified in every well used. The set-level comparison used GO cellular component sets with 15 to 500 detected members, without the generic containers, and of two sets sharing more than half of the detected members of the smaller kept the one with the larger absolute discovery effect (74 sets). A set’s donor value is the mean inclusion-over-host difference of its detected members minus that of all other proteins. Discovery donor values were tested against zero by one-sample t-test with BH correction across sets. Cohort means were compared by Pearson and Spearman correlation and sign agreement over all sets or single proteins (Figure S6m) and over discovery hits. Figures 4g, S6m and S6n come from the joint search, all other brain panels from a search of the discovery samples alone.

### Cross-system comparison

#### Signature and published proteomes

The three aggregate-against-host contrasts are those of Figure 5a. The brain contrast is a two-group limma fit in disease donors on one pseudobulk row per donor and group (the mean log2 intensity over that donor’s pools), over all sampled regions and without a brain-bank covariate, requiring at least three donors per group and a value in every donor row. Genes were matched by upper-case symbol, each system contributing its protein group with the highest detection fraction. A gene entered when quantified in at least 50% of a system’s samples (7,038 HEK293, 5,665 iDA, 4,703 brain) and given a fold change in all three contrasts. Shared up and shared down genes have log2FC ≥ 0.3 or ≤ −0.3 (set in advance) and FDR < 0.05 (added later) in every contrast, and the classification by fold change alone is also reported. The two-system signature of iDA neurons and brain (Figure 5b) was built in the same way from the genes detected in both. Agreement was measured by Pearson and Spearman correlation, with 10,000 label permutations for Pearson’s *r*. Published aggregate proteomes come from a workbook of 40 gene lists tiered by hand (Supplementary Table 4). The 21 lists tiered as primary aggregate composition or canonical histology were reduced to 18 independent units by merging three hand-specified duplicate pairs, and the consensus holds the genes in at least 3 of the 18. Symbols were matched in upper case, without alias or ortholog mapping. The Novel groups of Figures 5b and S7c hold shared-up genes absent from all 35 α-synuclein lists, which together name 2,842 genes. A published gene counted as detected in a system when it had a value in at least one aggregate sample (EA, somatic or neuritic aggregates, round or elongated inclusions) after that system’s completeness filter (Figure S7e). The brain-only enriched set (Figure S7f) holds the genes with log2FC > 0.3 in round inclusions against DAB-positive neurons or in elongated inclusions against neuropil.

#### Complexes and split assemblies

Complexes came from the Complex Portal (human, downloaded 10 September 2026). Participants were mapped to primary UniProt gene symbols after dropping nested complexes, small molecules and RNAs and expanding bracketed paralogue alternatives, which left 777 of 2,498 complexes with at least four distinct symbols. Complexes sharing more than 0.8 of their members (Jaccard index) were collapsed to the one with the most subunits. In each contrast a complex with at least four quantified subunits was analyzed. Its fold change is the median subunit log2FC, the standard deviation of its subunit fold changes was compared with the mean of 200 random gene sets of the same size, matched on abundance quartile and on a compartment class from UniProt subcellular-location keywords, and its direction was tested by one-sample t-test of subunit fold changes against zero. Gene Ontology pairs are two complex terms that are parts of the same protein-containing complex term and keep at least three genes each after removing shared genes. Protein Data Bank pairs are complexes without a shared subunit that each contribute at least two subunits to one structure, in at least two entries. Hand-written rules on the part names assigned each pair a tier and a machine, tier 1 meaning that both parts are documented together in one particle. In each contrast, pairs were removed when a part had fewer than three quantified subunits or the smallest attainable exact p-value was at least 0.05, and identical and near-duplicate pairs (Jaccard index above 0.8 on the union of both parts) were collapsed, keeping the lower tier, then the larger pair. The remaining pairs were tested by two-sided Mann-Whitney U test, and pairs confounded by shared subunits or by being one machine at two assembly stages were removed after testing. Figure 5e shows, for brain round inclusions and iDA somatic aggregates, one tier-1 pair per machine among the pairs significant in at least one of the two, chosen by the number of these contrasts in which it is significant, then by the number of PDB entries.

#### Modules, properties and curated panels

Each organelle module is a GO cellular component term selected by the authors with the help of an AI assistant (GO release 26 July 2026, human annotations of 21 May 2026), together with its part-of descendants. A module required at least four members quantified in at least 50% of the samples of both groups. In brain, each donor contributed one value, the mean module score of its inclusion pools minus that of its host pools, and module pairs were tested by an intercept-only limma over the five donors. Nine properties were computed per gene symbol, from the longest UniProt sequence after removing non-standard characters and sequences shorter than 25 residues. They are CamSol solubility (CamSol web server, averaged per symbol), mean Kyte-Doolittle hydrophobicity, net charge per residue ((K + R − D − E)/length, absolute in Figure 5f), fraction of charged residues, charge blockiness (mean absolute charge per residue over 25-residue windows minus the absolute whole-protein value), order propensity (fraction of W, C, F, I, Y, V, L and N), aromaticity (fraction of F, W and Y), log10 length and the number of transmembrane segments annotated in UniProt. Property correlations required at least 20 complexes and 200 proteins. Split-assembly differences between the mean properties of the two parts used each part’s members quantified in either contrast of the figure, at least four per part. The seven RNA-binding proteins (Figure S7d) were compared with the other nuclear genes of each contrast (GO:0005634 and its part-of descendants, detected in at least 50% of a system’s samples). Per gene, the log2FC was compared with the background median and with a local linear fit of background log2FC on log2 intensity (up to 300 nuclear genes within 1 log2 unit, at least 50), by empirical two-sided p-values with BH correction over the seven genes. As a set, the panel was tested by Mann-Whitney U test against the background and by a matched permutation of its median residual (10,000 draws, one residual from each gene’s window).

### Phosphoproteomics

#### Processing

Contaminant protein groups were first removed from the Spectronaut phosphopeptide and proteome reports. Blank wells and conditions with fewer than three wells were dropped. Wells with fewer Class I sites (localization probability ≥ 0.75 in that run) than 0.60 times the median of their condition were excluded from both reports (nine wells). Precursors were collapsed per protein group to modified peptides (sum per run) and to phosphosites (median over all precursors carrying the site, whose localization probability in a run is the highest among them), separately per multiplicity (phosphates per precursor, capped at three). Log2 values of 0 and 1 were set to missing. A site with a localization probability ≥ 0.75 in at least 50% of a condition’s replicates was kept in all of them, and otherwise its values below 0.75 were set to missing. Sites and modified peptides quantified in at least 70% of the replicates of at least one condition were kept. Within a condition, sites quantified in at least 50% of the replicates were imputed by k-nearest neighbors, the neighbors being the sites with the most similar profiles over all wells (*k* = max(3, ⌊√*n*⌋) for *n* wells). Sites detected in at most one replicate of a condition were drawn from a Gaussian distribution with a mean 1.8 standard deviations below the well’s mean and a width of 0.3 standard deviations, both from the well after kNN imputation (seed 42), and other missing values remained. The proteome (log2, zeros as missing) was filtered and imputed in the same way. Sites were normalized to their parent protein by subtracting its mean log2 intensity in the same condition, sites without a quantified parent were dropped, and observed and imputed matrices were both normalized. The phosphosite PCA used the imputed Class I sites with a value in every well, before normalization.

#### Testing and site presence

Class I sites were tested by limma on the normalized imputed matrix in 19 contrasts, the BafA1 effect within each fraction and, with and without BafA1, PFF against PBS in cells without aggregates and every pair of the four PFF fractions. Each contrast was fitted on its two groups and on the sites with measured values in at least 60%, and at least two, of the replicates of both groups. Sites detected in excised aggregates and not in rest cells (thresholds in Figure S9g, h) were counted per site and multiplicity on the observed matrix, and the multiplicity detected in most EA replicates was chosen as representative after selection. A site was constitutive when detected in at least 50% of the replicates of PBS cells without aggregates, PFF-induced when detected in at least 50% of those of PFF cells with or without aggregates, and aggregate-exclusive otherwise.

#### α-Synuclein C-terminus and its enzymes

SNCA phosphopeptides containing the C-terminal tryptic sequence (EMPSEE motif spanning Ser129) were summed in linear space over the observed peptide matrix, with unmeasured variants as zero, returned to log2 and normalized by subtracting the condition mean of log2 SNCA protein intensity. Acceptor residues were assigned from y ions in the fragment-level Spectronaut export. A y ion counted as phosphorylated when its calibrated m/z matched its calculated m/z, including methionine oxidation, plus 79.9663 Th divided by its charge (tolerance 0.02 Th). The y5, y8, y13 and y16 ions add Tyr136, Tyr133, Ser129 and Tyr125. Raw peak areas of phosphorylated y ions without neutral loss were summed per well over precursors and charge states and divided by the condition mean of linear SNCA protein intensity, and a condition’s value is log2 of the mean over its wells, with undetected wells as zero. Kinases and phosphatases linked to α-synuclein were grouped by enzyme class and target residue from a hand-assembled list. Their fold changes come from limma on the imputed proteome, and that of the C-terminal phosphopeptide from limma on the observed normalized peptide matrix. The confidence level of the intervals is one minus the geometric mean of the largest p-value rejected at FDR 0.05 and the smallest not rejected, with degrees of freedom recovered from the moderated *t* and its p-value.

## AI-use statement

The authors used Claude (Anthropic; Claude Opus 4.7, Opus 4.8, Opus 5 and Opus 5.5, accessed through Claude Code between April and October 2026) to write and debug code for image analysis, data analysis and figure plotting, and to draft and edit the manuscript text. The authors reviewed all code, checked all results and figures against the underlying data, revised all text and take responsibility for the content of this article. No image or figure panel was produced by an image-generation model.

## Data availability

The mass spectrometry proteomics data will become available upon publication.

## Ethics and consent

Post-mortem human brain tissue was obtained from two brain banks: the Neurobiobank München (NBM; Munich, Germany) and the Brain and Body Donation Program (BBDP) of the Banner Sun Health Research Institute (BSHRI; Sun City, Arizona, USA).

For tissue obtained from the NBM, written informed consent for tissue donation and research use was obtained from all donors during their lifetime or from their legal next of kin in accordance with the brain bank’s governing protocol. The collection, storage, and distribution of human post-mortem tissue by the NBM were approved by the Ethics Committee of the LMU Munich (#13-345).

For tissue obtained from the BBDP, all participants, or their legally authorized representative where a participant was unable to provide consent, gave written informed consent and HIPAA authorization for the donation of brain and bodily tissue and its use in research during their lifetime. The collection, storage, and distribution of human post-mortem tissue by the BBDP were approved by the Western Institutional Review Board (WCG IRB; protocol #20120821).

Use of the tissue for the present study was approved by the Ethics Committee of the LMU Munich (#23-0982).

All procedures were conducted in accordance with the Declaration of Helsinki.

## Author contributions

M.O. and M.M. conceived the study and designed experiments; M.O., S.V.G., A.L.M.M., C.S.S., V.R., D.O., U.D., J.P., T.H., E.U., K.M., Y.J., C.U., and Y.S. performed the experiments. M.O., L.D., A.S. and F.K. developed the image analysis. M.O., S.B.-M. and E.R. performed data analyses. M.O. and M.M. wrote the manuscript with input from all authors; M.M., F.U.H. and H.A.L. supervised the work and acquired funding. All authors read and approved the final manuscript.

## Code availability

All data-analysis code is based on the open-source alphapepttools package (https://github.com/MannLabs/alphapepttools). Full analysis pipelines, including preprocessing and figure generation, is available under https://github.com/MarcMarc94/aggregateDVP_analysis.

## Acknowledgments

The following scientific core facility at the Max Planck Institute of Biochemistry was used: Imaging Facility (RRID:SCR_025739). M.O. was supported by the HORIZON-MSCA-2023-PF-01-01 project HETLEWY, no. 101151819. A.L.M.M was supported by EPFL and fondation Bru. This project was also supported by the Michael J. Fox Foundation for Parkinson’s Research. We are grateful to the Banner Sun Health Research Institute Brain and Body Donation Program of Sun City, Arizona for the provision of human biological materials. The Brain and Body Donation Program has been supported by the NINDS (U24 NS072026), the NIA (P30 AG019610 and P30 AG072980), the Arizona Department of Health Services (contract 05700), the Arizona Biomedical Research Commission (contracts 4001, 0011, 05-901 and 1001) and the Michael J. Fox Foundation for Parkinson’s Research.

## References

1. Spillantini, M. G. et al. α-Synuclein in Lewy bodies. Nature 388, 839–840 (1997).

2. Glenner, G. G. & Wong, C. W. Alzheimer’s disease: Initial report of the purification and characterization of a novel cerebrovascular amyloid protein. Biochemical and Biophysical Research Communications 120, 885–890 (1984).

3. Masters, C. L., et al. Amyloid plaque core protein in Alzheimer disease and Down syndrome. Proc. Natl. Acad. Sci. U.S.A. 82, 4245–4249 (1985).

4. Kang, J. et al. The precursor of Alzheimer’s disease amyloid A4 protein resembles a cell-surface receptor. Nature 325, 733–736 (1987).

5. Grundke-Iqbal, I., et al. Abnormal phosphorylation of the microtubule-associated protein tau (tau) in Alzheimer cytoskeletal pathology. Proc. Natl. Acad. Sci. U.S.A. 83, 4913–4917 (1986).

6. Goedert, M., Wischik, C. M., Crowther, R. A., Walker, J. E. & Klug, A. Cloning and sequencing of the cDNA encoding a core protein of the paired helical filament of Alzheimer disease: identification as the microtubule-associated protein tau. Proc. Natl. Acad. Sci. U.S.A. 85, 4051–4055 (1988).

7. Neumann, M. et al. Ubiquitinated TDP-43 in Frontotemporal Lobar Degeneration and Amyotrophic Lateral Sclerosis. Science 314, 130–133 (2006).

8. Arai, T. et al. TDP-43 is a component of ubiquitin-positive tau-negative inclusions in frontotemporal lobar degeneration and amyotrophic lateral sclerosis. Biochemical and Biophysical Research Communications 351, 602–611 (2006).

9. Dorsey, E. R. et al. Global, regional, and national burden of Parkinson’s disease, 1990–2016: a systematic analysis for the Global Burden of Disease Study 2016. The Lancet Neurology 17, 939–953 (2018).

10. Li, M., Ye, X., Huang, Z., Ye, L. & Chen, C. Global burden of Parkinson’s disease from 1990 to 2021: a population-based study. BMJ Open 15, e095610 (2025).

11. Mastenbroek, S. E. et al. Disease progression modelling reveals heterogeneity in trajectories of Lewy-type α-synuclein pathology. Nat Commun 15, 5133 (2024).

12. Alafuzoff, I. et al. Staging/typing of Lewy body related α-synuclein pathology: a study of the BrainNet Europe Consortium. Acta Neuropathol 117, 635–652 (2009).

13. Shahmoradian, S. H. et al. Lewy pathology in Parkinson’s disease consists of crowded organelles and lipid membranes. Nat Neurosci 22, 1099–1109 (2019).

14. Lewis, A. J. et al. Ultrastructural diversity and subcellular organization of nigral Lewy pathology in Parkinson’s disease. Nat Commun 17, 7254 (2026).

15. Moors, T. E. et al. The subcellular arrangement of alpha-synuclein proteoforms in the Parkinson’s disease brain as revealed by multicolor STED microscopy. Acta Neuropathol 142, 423–448 (2021).

16. Kuzuhara, S., Mori, H., Izumiyama, N., Yoshimura, M. & Ihara, Y. Lewy bodies are ubiquitinated. Acta Neuropathol 75, 345–353 (1988).

17. Kuusisto, E., Parkkinen, L. & Alafuzoff, I. Morphogenesis of Lewy Bodies: Dissimilar Incorporation of α-Synuclein, Ubiquitin, and p62. J Neuropathol Exp Neurol 62, 1241–1253 (2003).

18. Goldman, J. E., Yen, S.-H., Chiu, F.-C. & Peress, N. S. Lewy Bodies of Parkinson’s Disease Contain Neurofilament Antigens. Science 221, 1082–1084 (1983).

19. Fujiwara, H. et al. α-Synuclein is phosphorylated in synucleinopathy lesions. Nat Cell Biol 4, 160–164 (2002).

20. Anderson, J. P. et al. Phosphorylation of Ser-129 Is the Dominant Pathological Modification of α-Synuclein in Familial and Sporadic Lewy Body Disease *. Journal of Biological Chemistry 281, 29739– 29752 (2006).

21. Mahul-Mellier, A.-L. et al. The process of Lewy body formation, rather than simply α-synuclein fibrillization, is one of the major drivers of neurodegeneration. Proceedings of the National Academy of Sciences 117, 4971–4982 (2020).

22. Leitão, A. D. G. et al. Selectivity of Lewy body protein interactions along the aggregation pathway of α-synuclein. Commun Biol 4, 1124 (2021).

23. Henderson, M. X. et al. Unbiased Proteomics of Early Lewy Body Formation Model Implicates Active Microtubule Affinity-Regulating Kinases (MARKs) in Synucleinopathies. J. Neurosci. 37, 5870–5884 (2017).

24. Bhattacharjee, P. et al. Mass Spectrometric Analysis of Lewy Body-Enriched α-Synuclein in Parkinson’s Disease. J. Proteome Res. 18, 2109–2120 (2019).

25. Cappelletti, V. et al. Dynamic 3D proteomes reveal protein functional alterations at high resolution in situ. Cell 184, 545–559.e22 (2021).

26. Killinger, B. A. et al. In situ proximity labeling identifies Lewy pathology molecular interactions in the human brain. Proceedings of the National Academy of Sciences 119, e2114405119 (2022).

27. Leverenz, J. B. et al. Proteomic Identification of Novel Proteins in Cortical Lewy Bodies. Brain Pathology 17, 139–145 (2007).

28. Method of the Year 2024: spatial proteomics. Nat Methods 21, 2195–2196 (2024).

29. Goltsev, Y. et al. Deep Profiling of Mouse Splenic Architecture with CODEX Multiplexed Imaging. Cell 174, 968–981.e15 (2018).

30. Giesen, C. et al. Highly multiplexed imaging of tumor tissues with subcellular resolution by mass cytometry. Nat Methods 11, 417–422 (2014).

31. Angelo, M. et al. Multiplexed ion beam imaging of human breast tumors. Nat Med 20, 436–442 (2014).

32. Ngai, Y. T., Lau, D., Mittal, P. & Hoffmann, P. Mini Review: Highlight of Recent Advances and Applications of MALDI Mass Spectrometry Imaging in 2024. Analytical Science Advances 6, e70016 (2025).

33. Guzman, U. H. et al. Ultra-fast label-free quantification and comprehensive proteome coverage with narrow-window data-independent acquisition. Nat Biotechnol 42, 1855–1866 (2024).

34. Makhmut, A., Qin, D., Hartlmayr, D., Seth, A. & Coscia, F. An Automated and Fast Sample Preparation Workflow for Laser Microdissection Guided Ultrasensitive Proteomics. Molecular & Cellular Proteomics 23, 100750 (2024).

35. Heymann, T. et al. Solid-Phase Extraction Capture (SPEC) in nanoliter volumes for fast, robust and ultra-sensitive proteomics. 2025.05.31.657165 Preprint at 10.1101/2025.05.31.657165 (2025).

36. Bubis, J. A. et al. Challenging the Astral mass analyzer to quantify up to 5,300 proteins per single cell at unseen accuracy to uncover cellular heterogeneity. Nat Methods 22, 510–519 (2025).

37. Mund, A. et al. Deep Visual Proteomics defines single-cell identity and heterogeneity. Nat Biotechnol 40, 1231–1240 (2022).

38. Rosenberger, F. A. et al. Spatial single-cell mass spectrometry defines zonation of the hepatocyte proteome. Nat Methods 20, 1530–1536 (2023).

39. Rosenberger, F. A. et al. Deep Visual Proteomics maps proteotoxicity in a genetic liver disease. Nature 642, 484–491 (2025).

40. Weiss, C. A. M. et al. Single-cell spatial proteomics maps human liver zonation patterns and their vulnerability to disruption in tissue architecture. Nat Metab 1–16 (2026) doi:10.1038/s42255-026-01459-2.

41. Foiani, M. S. et al. Spatial Single-Cell Proteomics Reveals Molecular Trajectories Of Tangle-Bearing Neurons In Alzheimer’s Disease. Preprint at 10.64898/2026.04.26.720932 (2026).

42. Mahul-Mellier, A.-L. et al. Modeling Parkinson’s pathology in human iPSC dopaminergic neurons uncovers key mechanisms of Lewy body formation and heterogeneity. Sci. Adv. 12, eaed8851 (2026).

43. Hasegawa, M. et al. Phosphorylated α-Synuclein Is Ubiquitinated in α-Synucleinopathy Lesions*. Journal of Biological Chemistry 277, 49071–49076 (2002).

44. Pachitariu, M. & Stringer, C. Cellpose 2.0: how to train your own model. Nat Methods 19, 1634–1641 (2022).

45. He, J. et al. Immunohistochemical staining, laser capture microdissection, and filter-aided sample preparation-assisted proteomic analysis of target cell populations within tissue samples. ELECTROPHORESIS 34, 1627–1636 (2013).

46. Huang, P. et al. Spatial proteome profiling by immunohistochemistry-based laser capture microdissection and data-independent acquisition proteomics. Analytica Chimica Acta 1127, 140–148 (2020).

47. Tangrea, M. A. et al. Immunoguided Microdissection Techniques. in Laser Capture Microdissection: Methods and Protocols (ed. Murray, G. I.) 57–66 (Humana Press, Totowa, NJ, 2011). doi:10.1007/978-1-61779-163-5_4.

48. Webb, J. L., Ravikumar, B., Atkins, J., Skepper, J. N. & Rubinsztein, D. C. α-Synuclein Is Degraded by Both Autophagy and the Proteasome *. Journal of Biological Chemistry 278, 25009–25013 (2003).

49. Winslow, A. R. et al. α-Synuclein impairs macroautophagy: implications for Parkinson’s disease. J Cell Biol 190, 1023–1037 (2010).

50. Chang, K.-H. & Chen, C.-M. The Role of Oxidative Stress in Parkinson’s Disease. Antioxidants 9, 597 (2020).

51. Mancias, J. D., Wang, X., Gygi, S. P., Harper, J. W. & Kimmelman, A. C. Quantitative proteomics identifies NCOA4 as the cargo receptor mediating ferritinophagy. Nature 509, 105–109 (2014).

52. Tian, Y. et al. FTH1 Inhibits Ferroptosis Through Ferritinophagy in the 6-OHDA Model of Parkinson’s Disease. Neurotherapeutics 17, 1796–1812 (2020).

53. Wu, K.-M. et al. Neuronal FAM171A2 mediates α-synuclein fibril uptake and drives Parkinson’s disease. Science 387, 892–900 (2025).

54. Sormanni, P., Aprile, F. A. & Vendruscolo, M. The CamSol method of rational design of protein mutants with enhanced solubility. Journal of Molecular Biology 427, 478–490 (2015).

55. Oeller, M. et al. Sequence-based prediction of the intrinsic solubility of peptides containing non-natural amino acids. Nat Commun 14, 7475 (2023).

56. Oliinyk, D. et al. nanoPhos enables ultra-sensitive and cell-type resolved spatial phosphoproteomics. 2025.05.29.656770 Preprint at 10.1101/2025.05.29.656770 (2025).

57. Burré, J. et al. α-Synuclein Promotes SNARE-Complex Assembly in Vivo and in Vitro. Science 329, 1663– 1667 (2010).

58. Diao, J. et al. Native α-synuclein induces clustering of synaptic-vesicle mimics via binding to phospholipids and synaptobrevin-2/VAMP2. eLife 2, e00592 (2013).

59. Olzscha, H. et al. Amyloid-like aggregates sequester numerous metastable proteins with essential cellular functions. Cell 144, 67–78 (2011).

60. Hosp, F. et al. Spatiotemporal Proteomic Profiling of Huntington’s Disease Inclusions Reveals Widespread Loss of Protein Function. Cell Reports 21, 2291–2303 (2017).

61. Tahmaz, I., Shahmoradi Ghahe, S. & Topf, U. Prefoldin Function in Cellular Protein Homeostasis and Human Diseases. Front. Cell Dev. Biol. 9, (2022).

62. Sitron, C. S. et al. α-Synuclein aggregates inhibit ESCRT-III through sequestration and collateral degradation. Molecular Cell 85, 3505–3523.e17 (2025).

63. Ghanem, S. S. et al. α-Synuclein phosphorylation at serine 129 occurs after initial protein deposition and inhibits seeded fibril formation and toxicity. Proceedings of the National Academy of Sciences 119, e2109617119 (2022).

64. Kumar, S. T., Donzelli, S., Chiki, A., Syed, M. M. K. & Lashuel, H. A. A simple, versatile and robust centrifugation-based filtration protocol for the isolation and quantification of α-synuclein monomers, oligomers and fibrils: Towards improving experimental reproducibility in α-synuclein research. Journal of Neurochemistry 153, 103–119 (2020).

65. Sheta, R., Teixeira, M., Idi, W. & Oueslati, A. Optimized protocol for the generation of functional human induced-pluripotent-stem-cell-derived dopaminergic neurons. STAR Protocols 4, 102486 (2023).

66. Mädler, S. C. et al. scPortrait integrates single-cell images into multimodal modeling. 2025.09.22.677590 Preprint at 10.1101/2025.09.22.677590 (2025).

67. Stirling, D. R. et al. CellProfiler 4: improvements in speed, utility and usability. BMC Bioinformatics 22, 433 (2021).

68. Marconato, L. et al. SpatialData: an open and universal data framework for spatial omics. Nat Methods 1–5 (2024) doi:10.1038/s41592-024-02212-x.

69. Ruifrok, A. C. & Johnston, D. A. Quantification of histochemical staining by color deconvolution. Anal Quant Cytol Histol 23, 291–299 (2001).

70. van der Walt, S. et al. scikit-image: image processing in Python. PeerJ 2, e453 (2014).

71. Skowronek, P. et al. Rapid and In-Depth Coverage of the (Phospho-)Proteome With Deep Libraries and Optimal Window Design for dia-PASEF. Molecular & Cellular Proteomics 21, 100279 (2022).

72. Demichev, V., Messner, C. B., Vernardis, S. I., Lilley, K. S. & Ralser, M. DIA-NN: neural networks and interference correction enable deep proteome coverage in high throughput. Nat Methods 17, 41–44 (2020).

73. Kistner, F., Grossmann, J. L., Sinn, L. R. & Demichev, V. QuantUMS: uncertainty minimisation enables confident quantification in proteomics. 2023.06.20.545604 Preprint at 10.1101/2023.06.20.545604 (2023).

74. Brennsteiner, V. et al. AlphaPeptTools: scverse-native analysis of mass spectrometry-based proteomics. 2026.09.11.750886 Preprint at 10.64898/2026.09.11.750886 (2026).

