## Extended Figures for "Aggregate Deep Visual Proteomics reveals selective organization of α-synuclein inclusions"

### Extended Data for: Aggregate Deep Visual Proteomics reveals selective organization of α-synuclein inclusions (Figures S1-10)

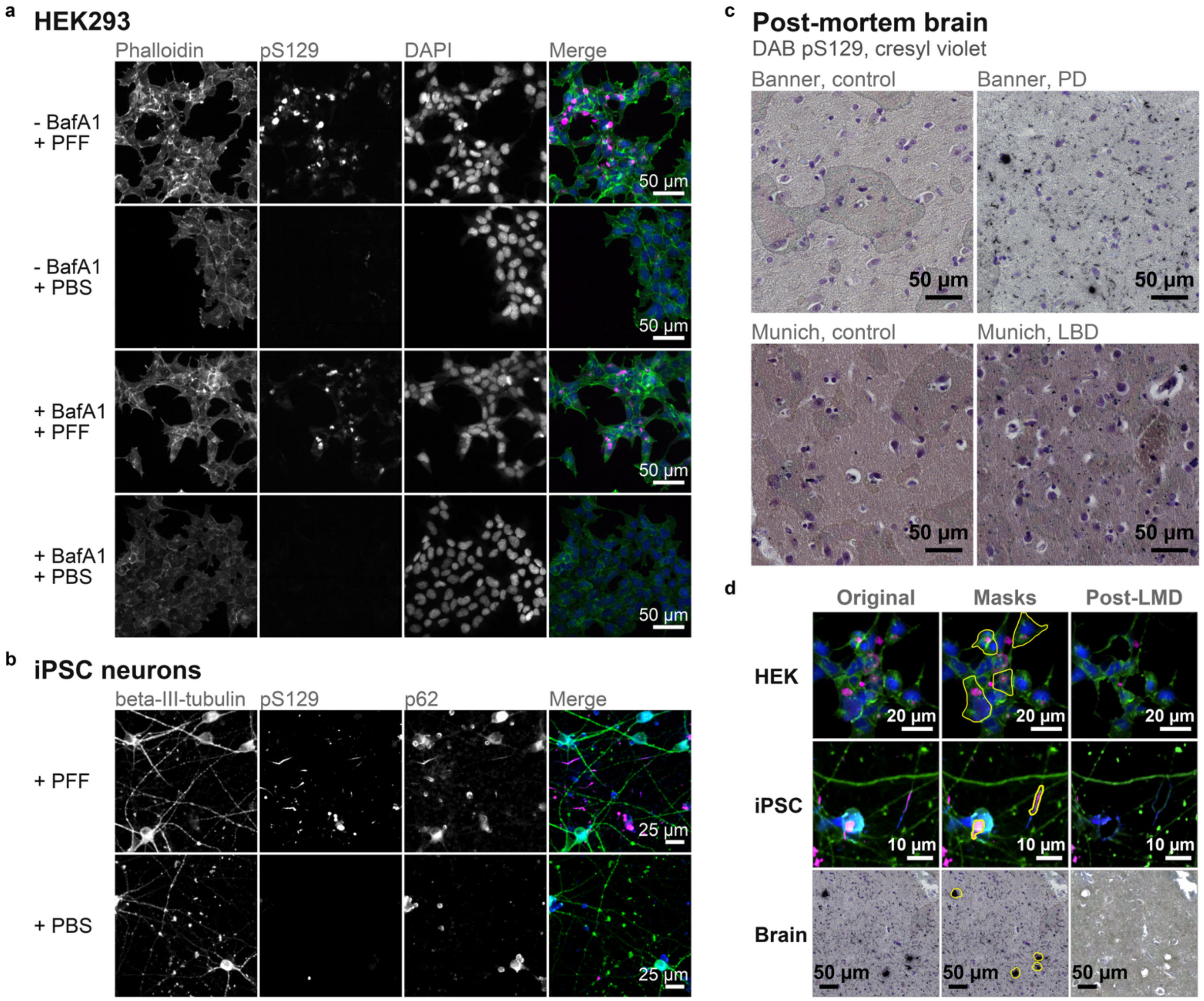


***Figure S1. Aggregate staining and laser capture in all three systems.***

*(a) HEK293 cells, four conditions (± bafilomycin A1, ± PFF): phalloidin, pS129 and DAPI as single greyscale channels and merged (phalloidin green, pS129 magenta, DAPI blue). (b) iPSC-derived neurons, + PFF and + PBS: beta-III-tubulin, pS129 and p62, merged in the same colour scheme. (c) Post-mortem middle temporal gyrus (Banner) and frontobasal cortex (Munich), DAB against pS129 with cresyl violet counterstain, one field per biobank and condition. (d) Cutting efficiency per system: the field before cutting, the same field with the capture outlines, and the field after laser microdissection. Within each system, pS129 is displayed on one intensity range shared by all conditions, its lower bound set from the unseeded condition and its upper bound from the seeded one. Beta-III-tubulin in (b) is scaled within each condition, as the PBS slides have approximately 2.5-fold lower background. Brightfield in (c) is unadjusted. The post-LMD brain scan in (d) was underexposed approximately 2.4-fold and was linearly rescaled per channel to the in-tissue median and spread of the pre-cutting scan. Magnification differs between blocks; scale bars are shown in each panel.*

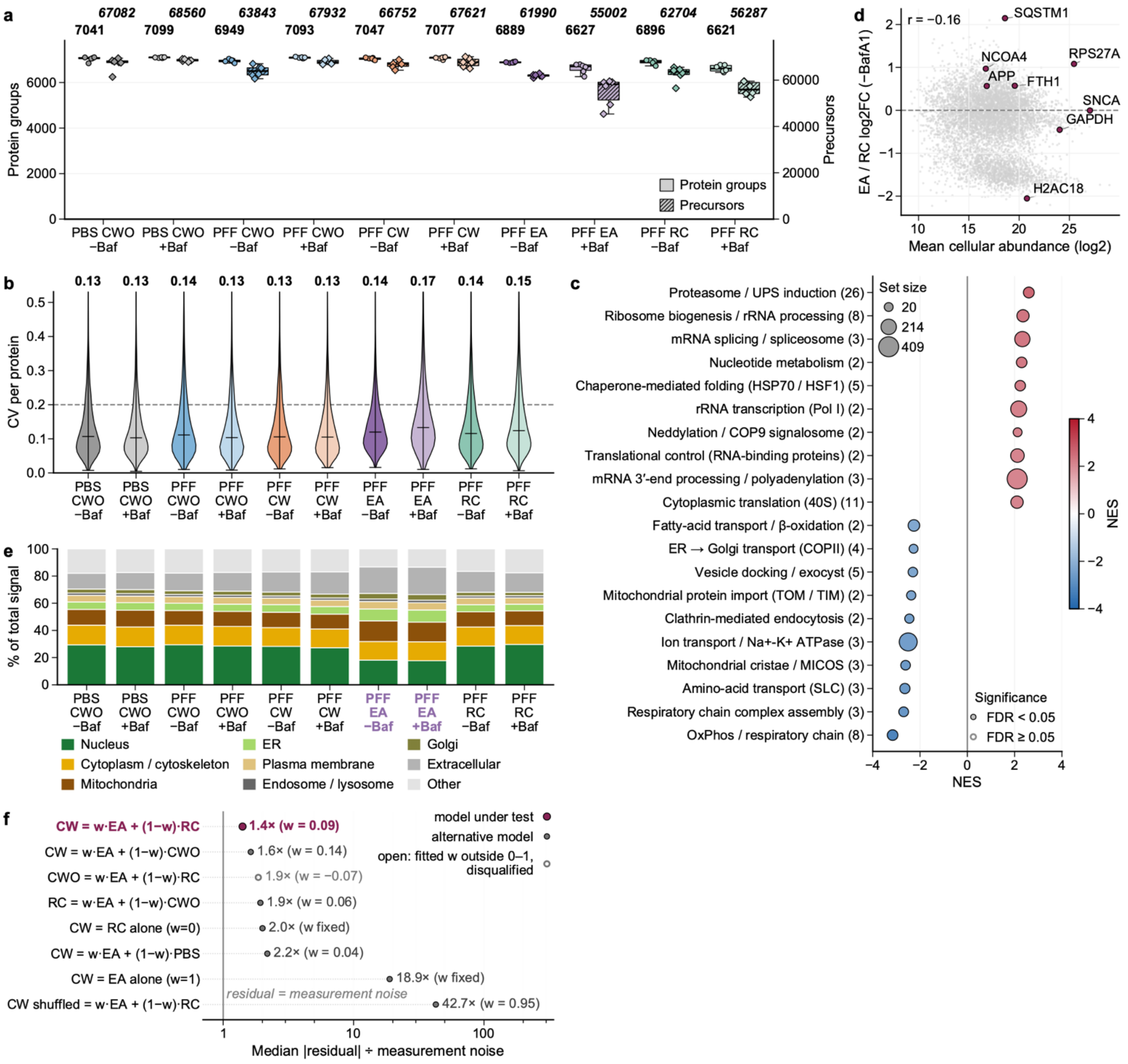


***Figure S2. Data quality, enrichment specificity and fraction composition in the HEK293 dataset.***
*(a) Protein groups (left axis) and precursors (right axis) per condition, n = 6 replicates. (b) Coefficient of variation per condition; means annotated, dashed line at 20%. (c) Gene set enrichment for PFF versus PBS in aggregate-free cells (−BafA1); positive normalised enrichment score (NES) indicates higher abundance after PFF. Related terms were collapsed into the categories shown, with the number of contributing terms in parentheses, dot area scaled to set size, and all categories at FDR < 0.05. (d) Mean cellular abundance (log2, averaged across cellular fractions) against aggregate enrichment (excised aggregate over residual cell body, EA / RC, −BafA1) for all quantified proteins, with representative proteins labelled; the weak Pearson correlation (r = −0.16) shows that enrichment is not driven by co-purification of abundant proteins. (e) Percentage of total protein signal assigned to each subcellular compartment across all ten conditions. (f) Median absolute per-protein residual divided by the bootstrap measurement noise for each mixing model, so 1 (grey line) means the residual matches noise. Dark red is the model under test, mixing the excised aggregate with its matched residual cell body; the other rows swap a component or the target for an unmatched fraction, fix the mixing weight w, or shuffle protein identities. Open symbols mark a fitted w outside 0 to 1, which disqualifies that row. Markers are the 1,113 proteins that differ between EA and CWO by the rule of Figure 2d, among the 5,700 proteins quantified in all 30 −BafA1 wells; rows ordered by residual.*

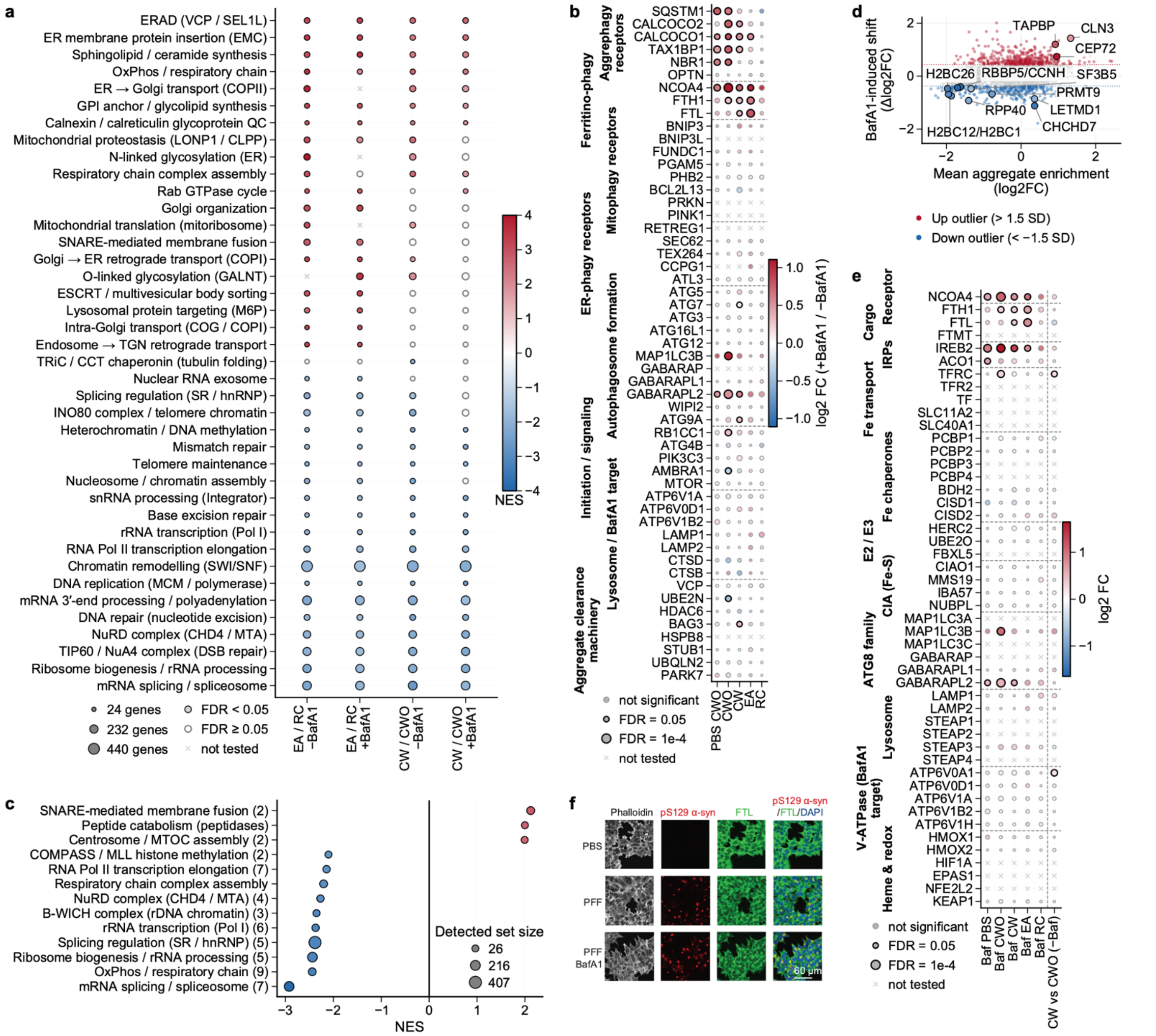


***Figure S3 . Pathway composition, autophagic flux and iron handling in HEK293 fractions under lysosomal blockade.*** *(a) Gene set enrichment for the excised aggregate against its residual cell body (EA / RC) and aggregate-bearing against aggregate-free cells (CW / CWO), each ± bafilomycin A1. Terms were collapsed; the top 20 and bottom 20 by maximum |NES| are shown. FDR is controlled within contrast, so admission on one of four is a display rule. (b) Bafilomycin A1 effect, log2 FC (+BafA1 / −BafA1), for autophagy receptors and core machinery by cargo route, per fraction (PBS, PFF). Positive values indicate accumulation under blockade, therefore degradation. (c) As (a), for the BafA1-induced shift: log2 FC (EA / RC, +BafA1) − log2 FC (EA / RC, −BafA1). Terms (FDR < 0.05) collapsed at a leading-edge overlap coefficient of 0.5; 13 of 26 clusters shown: all three positive, the ten most negative. (d) The shift per protein against mean aggregate enrichment, coloured beyond 1.5 SD; labels mark one representative protein per category in (c), not the largest shifts. (e) As (b), for 48 iron-handling proteins by functional category, across five fraction comparisons and bulk CW / CWO. Seventeen of them, FBXL5 among them, were not quantified in any comparison (×). (f) Immunofluorescence of HEK293 α-synuclein cells, PFF two days, then 100 nM BafA1 6 h where indicated; actin (phalloidin), pS129 α-synuclein, ferritin light chain (FTL). Colour: NES in (a, c), log2 FC in (b, e). Dot area: set size in (a, c), −log10 FDR in (b, e). In (a), filled dots pass FDR < 0.05; in (b, e), a black outline marks FDR < 0.05; × marks data not quantified.*

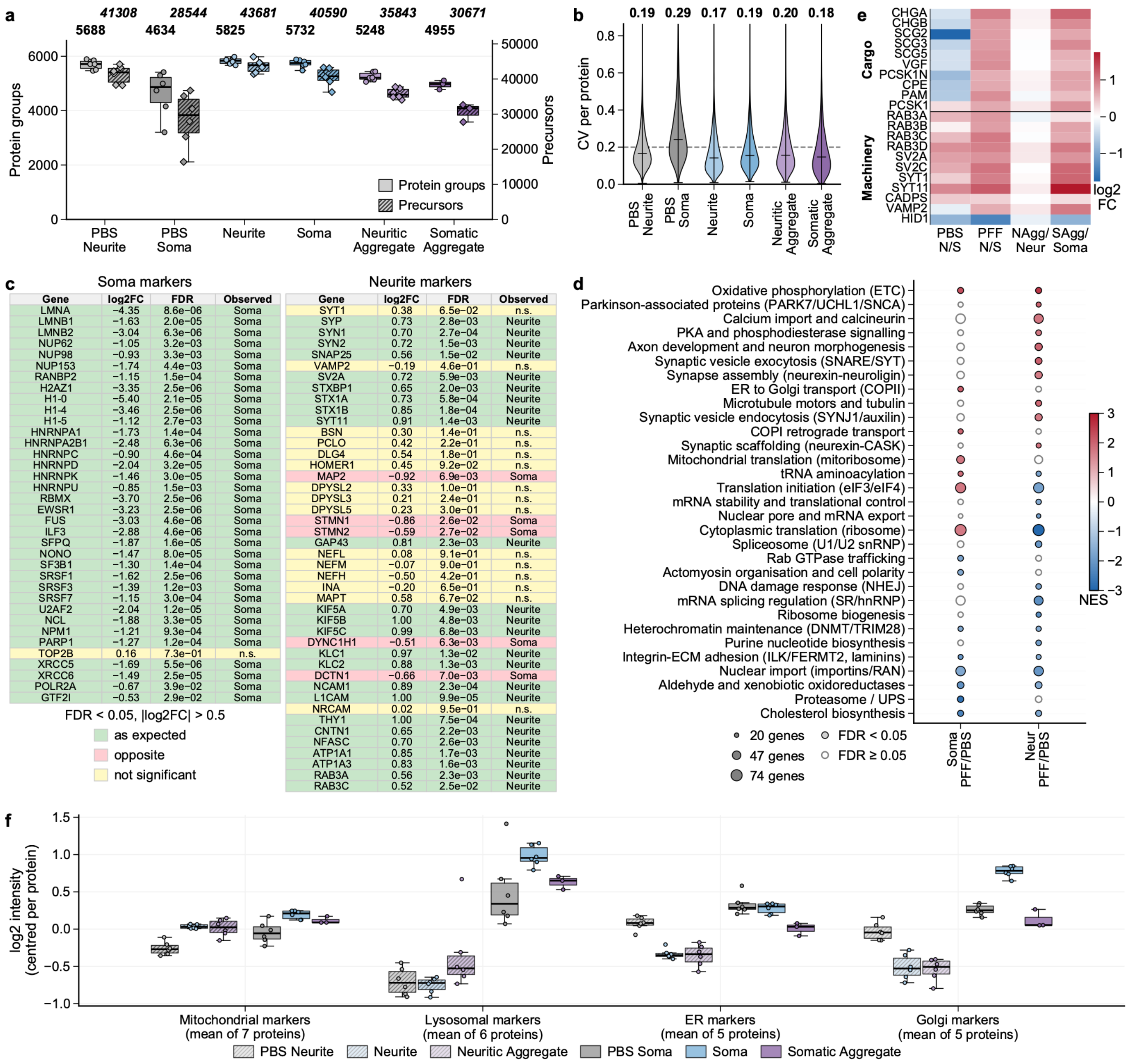


Soma markers

| Gene | log2FC | FDR | Observed |
|---|---|---|---|
| LMNA | −4.35 | 8.6e−06 | Soma |
| LMNB1 | −1.63 | 2.0e−05 | Soma |
| LMNB2 | −3.04 | 6.3e−06 | Soma |
| NUP62 | −1.05 | 3.2e−03 | Soma |
| NUP98 | −0.93 | 3.3e−03 | Soma |
| NUP153 | −1.74 | 4.4e−03 | Soma |
| RANBP2 | −1.15 | 1.5e−04 | Soma |
| H2AZ1 | −3.35 | 2.5e−06 | Soma |
| H1-0 | −5.40 | 2.1e−05 | Soma |
| H1-4 | −3.46 | 2.5e−06 | Soma |
| H1-5 | −1.12 | 2.7e−03 | Soma |
| HNRNPA1 | −1.73 | 1.4e−04 | Soma |
| HNRNPA2B1 | −2.48 | 6.3e−06 | Soma |
| HNRNPC | −0.90 | 4.6e−04 | Soma |
| HNRNPD | −2.04 | 3.2e−05 | Soma |
| HNRNPK | −1.46 | 3.0e−05 | Soma |
| HNRNPU | −0.85 | 1.5e−03 | Soma |
| RBMX | −3.70 | 2.5e−06 | Soma |
| EWSR1 | −3.23 | 2.5e−06 | Soma |
| FUS | −3.03 | 4.6e−06 | Soma |
| ILF3 | −2.88 | 4.6e−06 | Soma |
| SFPQ | −1.87 | 1.6e−05 | Soma |
| NONO | −1.47 | 8.0e−05 | Soma |
| SF3B1 | −1.30 | 1.4e−04 | Soma |
| SRSF1 | −1.62 | 2.5e−06 | Soma |
| SRSF3 | −1.39 | 1.2e−03 | Soma |
| SRSF7 | −1.15 | 3.0e−04 | Soma |
| U2AF2 | −2.04 | 1.2e−05 | Soma |
| NCL | −1.88 | 3.3e−05 | Soma |
| NPM1 | −1.21 | 9.3e−04 | Soma |
| PARP1 | −1.27 | 1.2e−04 | Soma |
| TOP2B | 0.16 | 7.3e−01 | n.s. |
| XRCC5 | −1.69 | 5.5e−06 | Soma |
| XRCC6 | −1.49 | 2.5e−05 | Soma |
| POLR2A | −0.67 | 3.9e−02 | Soma |
| GTF2I | −0.53 | 2.9e−02 | Soma |

Neurite markers

| Gene | log2FC | FDR | Observed |
|---|---|---|---|
| SYT1 | 0.38 | 6.5e−02 | n.s. |
| SYP | 0.73 | 2.8e−03 | Neurite |
| SYN1 | 0.70 | 2.7e−04 | Neurite |
| SYN2 | 0.72 | 1.5e−03 | Neurite |
| SNAP25 | 0.56 | 1.5e−02 | Neurite |
| VAMP2 | −0.19 | 4.6e−01 | n.s. |
| SV2A | 0.72 | 5.9e−03 | Neurite |
| STXBP1 | 0.65 | 2.0e−03 | Neurite |
| STX1A | 0.73 | 5.8e−04 | Neurite |
| STX1B | 0.85 | 1.8e−04 | Neurite |
| SYT11 | 0.91 | 1.4e−03 | Neurite |
| BSN | 0.30 | 1.4e−01 | n.s. |
| PCLO | 0.42 | 2.2e−01 | n.s. |
| DLG4 | 0.54 | 1.8e−01 | n.s. |
| HOMER1 | 0.45 | 9.2e−02 | n.s. |
| MAP2 | −0.92 | 6.9e−03 | Soma |
| DPYSL2 | 0.33 | 1.0e−01 | n.s. |
| DPYSL3 | 0.21 | 2.4e−01 | n.s. |
| DPYSL5 | 0.23 | 3.0e−01 | n.s. |
| STMN1 | −0.86 | 2.6e−02 | Soma |
| STMN2 | −0.59 | 2.7e−02 | Soma |
| GAP43 | 0.81 | 2.3e−03 | Neurite |
| NEFL | 0.08 | 9.1e−01 | n.s. |
| NEFM | −0.07 | 9.0e−01 | n.s. |
| NEFH | −0.50 | 4.2e−01 | n.s. |
| INA | −0.20 | 6.5e−01 | n.s. |
| MAPT | 0.58 | 6.7e−02 | n.s. |
| KIF5A | 0.70 | 4.9e−03 | Neurite |
| KIF5B | 1.00 | 4.8e−03 | Neurite |
| KIF5C | 0.99 | 6.8e−03 | Neurite |
| DYNC1H1 | −0.51 | 6.3e−03 | Soma |
| KLC1 | 0.97 | 1.3e−02 | Neurite |
| KLC2 | 0.88 | 1.3e−03 | Neurite |
| DCTN1 | −0.66 | 7.0e−03 | Soma |
| NCAM1 | 0.89 | 2.3e−04 | Neurite |
| L1CAM | 1.00 | 9.9e−05 | Neurite |
| NRCAM | 0.02 | 9.5e−01 | n.s. |
| THY1 | 1.00 | 7.5e−04 | Neurite |
| CNTN1 | 0.65 | 2.2e−03 | Neurite |
| NFASC | 0.70 | 2.6e−03 | Neurite |
| ATP1A1 | 0.85 | 1.7e−03 | Neurite |
| ATP1A3 | 0.83 | 1.6e−03 | Neurite |
| RAB3A | 0.56 | 2.3e−03 | Neurite |
| RAB3C | 0.52 | 2.5e−02 | Neurite |

***Figure S4. Quality, compartment identity and composition of the iPSC-derived dopaminergic neuron fractions.***

*(a) Protein groups (left axis) and precursors (right axis) per condition; n = 6, somatic aggregates n = 3. (b) Coefficient of variation per condition, means annotated, dashed line at 20%. (c) Log2 FC (neurite / soma) and FDR for 36 soma and 44 neurite markers in the PBS condition, with each marker's expected compartment and whether the observation matched. (d) Gene set enrichment for PFF versus PBS in aggregate-free soma and neurite regions of day-56 neurons (n = 6). The 2,467 protein groups quantified in every sample were ranked by the Welch t-statistic, without imputation, against Reactome and GO Biological Process. Significant sets were clustered by gene overlap and curated; the 31 rows are the modules with the largest positive and negative NES (Methods). (e) Log2 FC for dense-core-vesicle cargo and machinery in four comparisons: PBS and PFF neurites over soma, neuritic aggregates over PFF neurites, somatic aggregates over PFF soma. (f) Organelle marker intensity per sample. Each marker was centred on its own mean across samples, then averaged within each family, which puts the families on one scale and preserves all differences between conditions: a family depleted from an aggregate would sit below its source compartment. Boxes show the median and interquartile range, whiskers 1.5 × IQR, points replicates; hatching marks neurites. In (d), colour gives NES (red higher in PFF), dot size the number of detected proteins, and filled dots pass FDR < 0.05.*

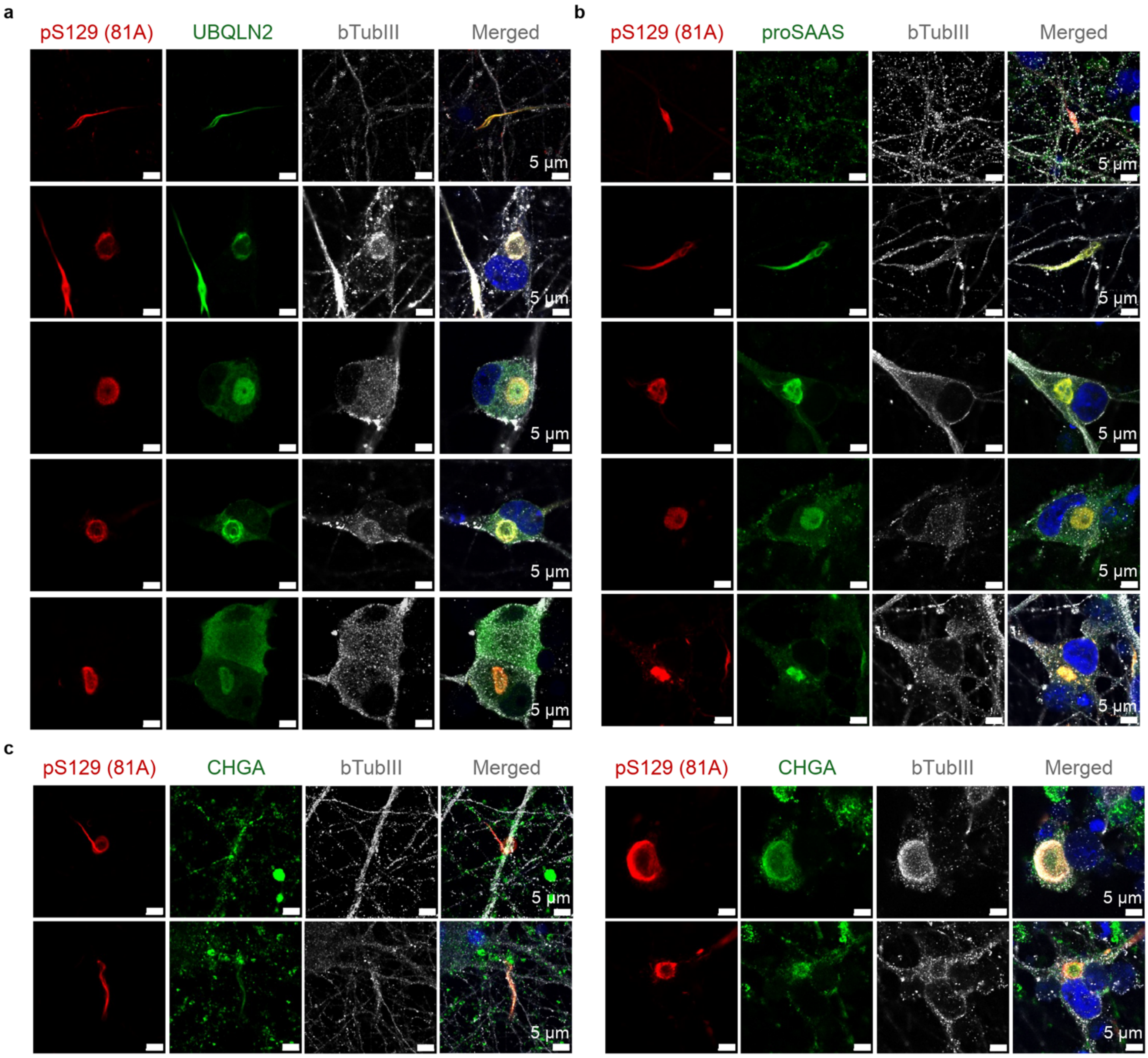


***Figure S5. Immunofluorescence of inclusion-associated proteins in iPSC-derived dopaminergic neurons.***
*(a–c) Neurons 56 days after PFF seeding, stained for UBQLN2 (a), proSAAS/PCSK1N (b) or chromogranin A (CHGA) (c), with somatic and neuritic inclusions shown for each target. (a) UBQLN2 co-localises with pS129 in both inclusion types. (b, c) Some neuritic inclusions are positive for proSAAS or CHGA, but most pS129-positive neurites are negative for both (top rows). Co-localising examples are shown in the rows beneath. Target protein, green; pS129 α-synuclein, red; beta-III-tubulin, grey; DAPI, blue. Scale bars, 5 µm. For completeness reasons, the same merged images that are shown in Figure 3 are reproduced here as well.*

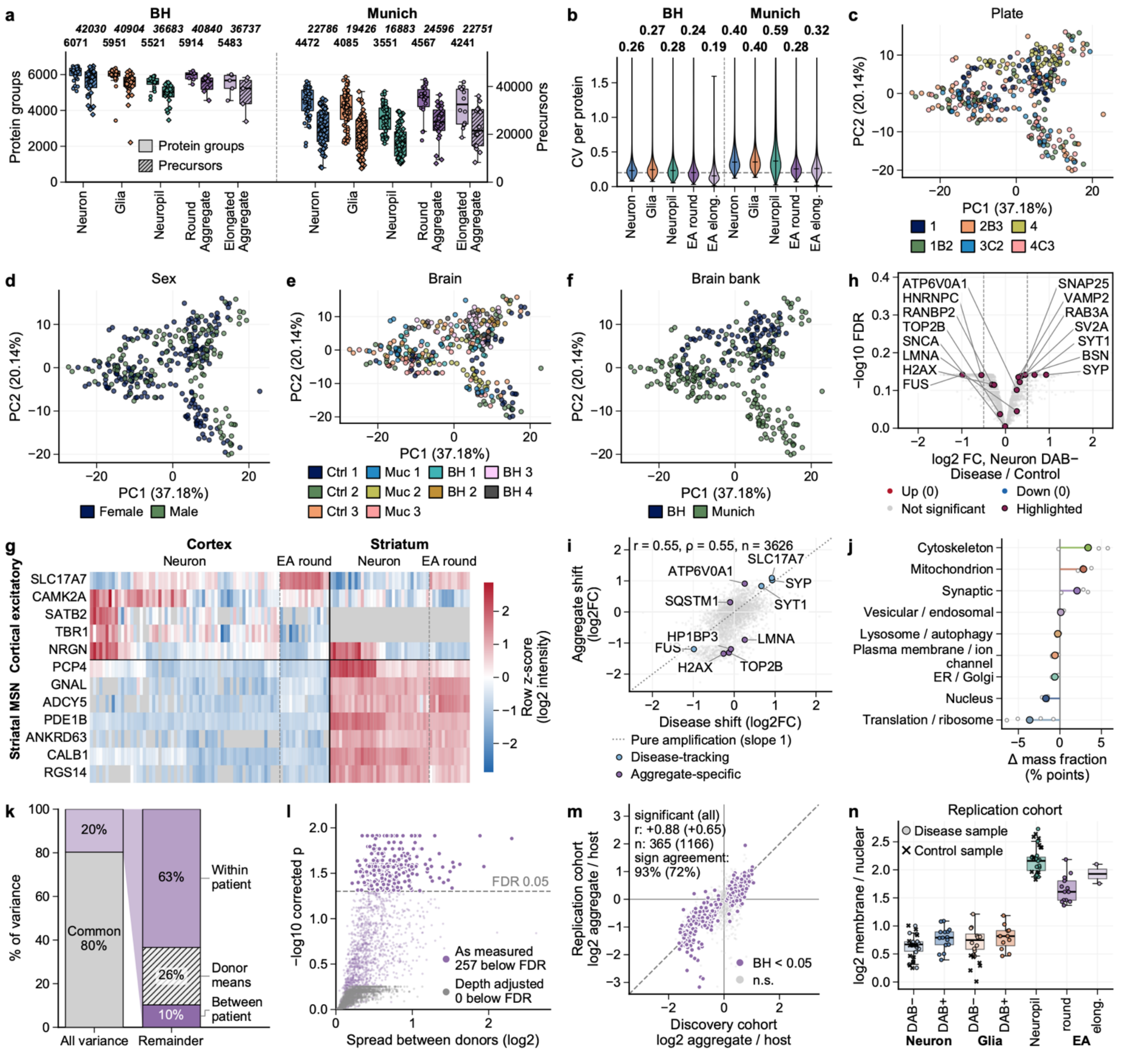


*Figure S6. Data quality, technical controls and validation of the post-mortem brain dataset.*

*(a, b) Protein groups and precursors per sample (a) and coefficient of variation (b) per cell type, by brain bank (Banner Health and Munich; 332 samples); means annotated. (c–f) Principal component analysis of all 332 samples on the 802 protein groups quantified in every one, one projection coloured by plate (c), sex (d), donor (e) and brain bank (f). None separates the samples. (g) Row z-scored intensity for cortical excitatory and striatal medium-spiny-neuron markers by region and fraction; round aggregates follow their host region. Grey, not detected. (h) Aggregate-free neurons from diseased against control donors (limma moderated t on donor means, n = 3 versus 5, BH-FDR), with Figure 4d aggregate markers overlaid. (i) Aggregate shift (round versus DAB+ neuron, n = 5 paired donors) against that disease shift; r = 0.55. Blue, disease-tracking; violet, aggregate-specific. (j) Paired within-donor change in compartment mass fraction, percentage points (n = 4 cortical disease donors); lollipops, mean; circles, donors. (k) Variance in the pool-level aggregate-over-host ratio, sum-of-squares units; the second bar subdivides the remainder of the first, the between-patient term corrected for sampling error in each donor mean. (l) Between-patient difference per protein before and after removing the linear effect of identification count; exact permutation over all 72,072 assignments, BH across proteins. (m) Donor-level aggregate-over-host ratio per protein, discovery against replication cohort (different brains and batch, searched jointly). Violet, discovery FDR < 0.05; grey, all other proteins; dashed, identity. With three replication donors, agreement is judged on effect size and direction. (n) Summed membrane and vesicular over summed nuclear and translational intensity per sample in the replication cohort; coloured dots and boxes, disease; black crosses, control. Cohorts differ in absolute level, so fractions are compared within a cohort. Boxes, median and interquartile range; whiskers, 1.5 × IQR.*

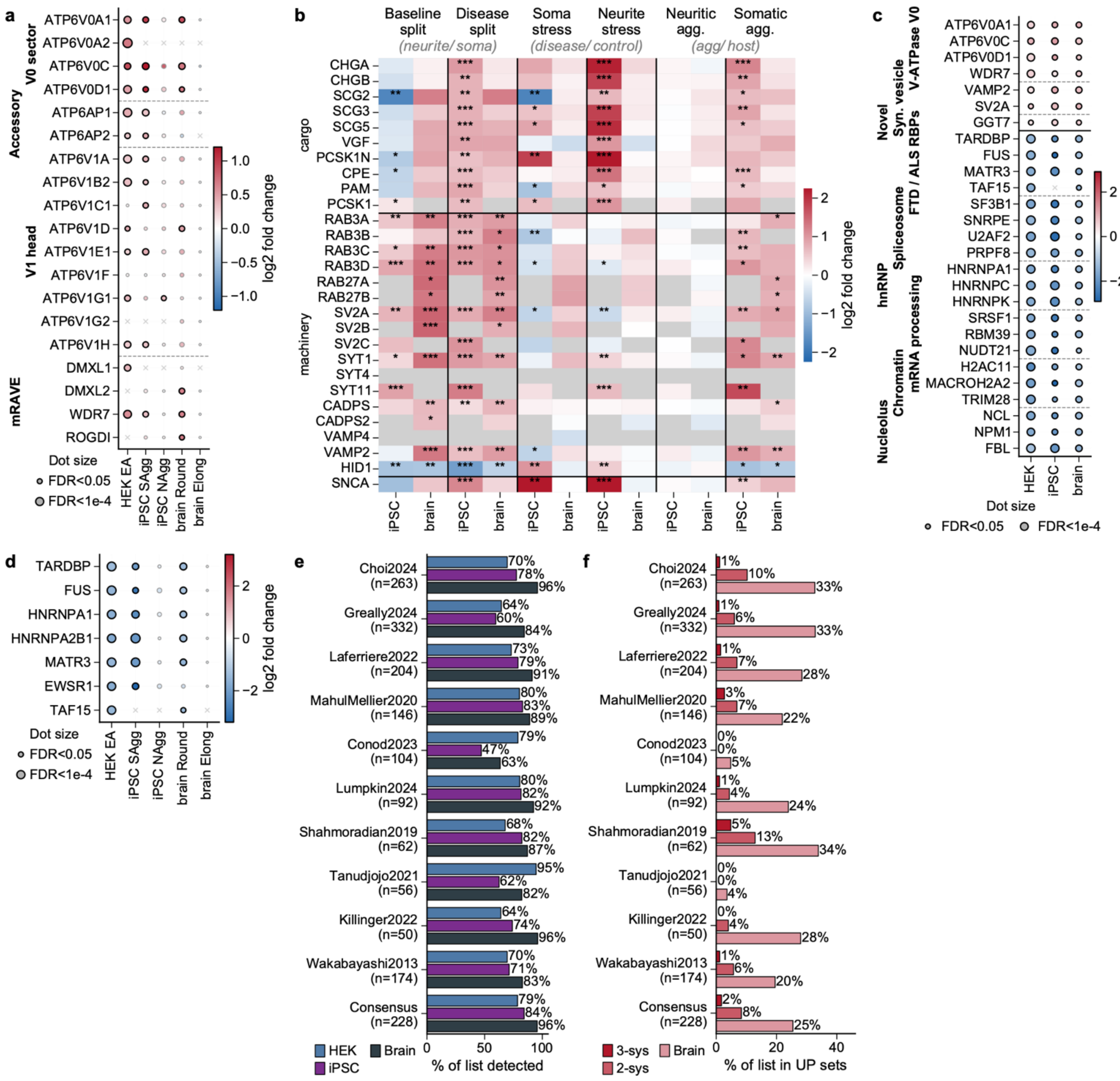

*Figure S7. Cross-system aggregate composition and overlap with published α-synuclein datasets.*
*(a) V-ATPase subunits and the mRAVE assembly factors by sector across five aggregate-versus-host contrasts: HEK293 EA versus RC, iPSC somatic and neuritic aggregates versus soma and neurites, brain round aggregates versus DAB+ neurons and elongated aggregates versus neuropil. Black outlines mark FDR < 0.05. ATP6V0A1, ATP6V0C, ATP6V0D1 and WDR7 are the four proteins of the panel enriched in all three systems (c). (b) Dense-core-vesicle cargo, release machinery and SNCA across six cross-system comparisons, the iPSC and brain columns adjacent within each. HEK293 is omitted because it does not express DCV cargo. Asterisks, BH-FDR (* < 0.05, ** < 0.01, *** < 0.001). (c) The stringent three-system signature: proteins at |log2 FC| ≥ 0.3 and FDR < 0.05 with concordant sign in HEK EA versus RC, iPSC somatic aggregate versus PFF soma, and brain round aggregate versus DAB+ neuron. All 7 universal-UP proteins are shown with 20 of the 256 universal-DOWN, grouped by class. (d) Seven ALS/FTD aggregation-prone RNA-binding proteins (TARDBP, FUS, HNRNPA1, HNRNPA2B1, MATR3, EWSR1, TAF15) across the five aggregate-versus-host contrasts of (a). They are depleted from every somatic aggregate fraction. (e) Percentage of each published α-synuclein list detected in the aggregate fractions of each system, for ten datasets and their consensus (members of at least three studies). (f) Percentage of each list that is enriched in the aggregate, by membership of three UP sets: brain (log2 FC > 0.3 in round aggregates against DAB+ neurons or in elongated aggregates against neuropil, without an FDR threshold), two-system and three-system universal-UP, counts in parentheses; the enriched fraction falls as cross-system stringency tightens. Colour gives log2 fold change and dot size −log10 FDR throughout; × marks proteins not quantified in that system.*

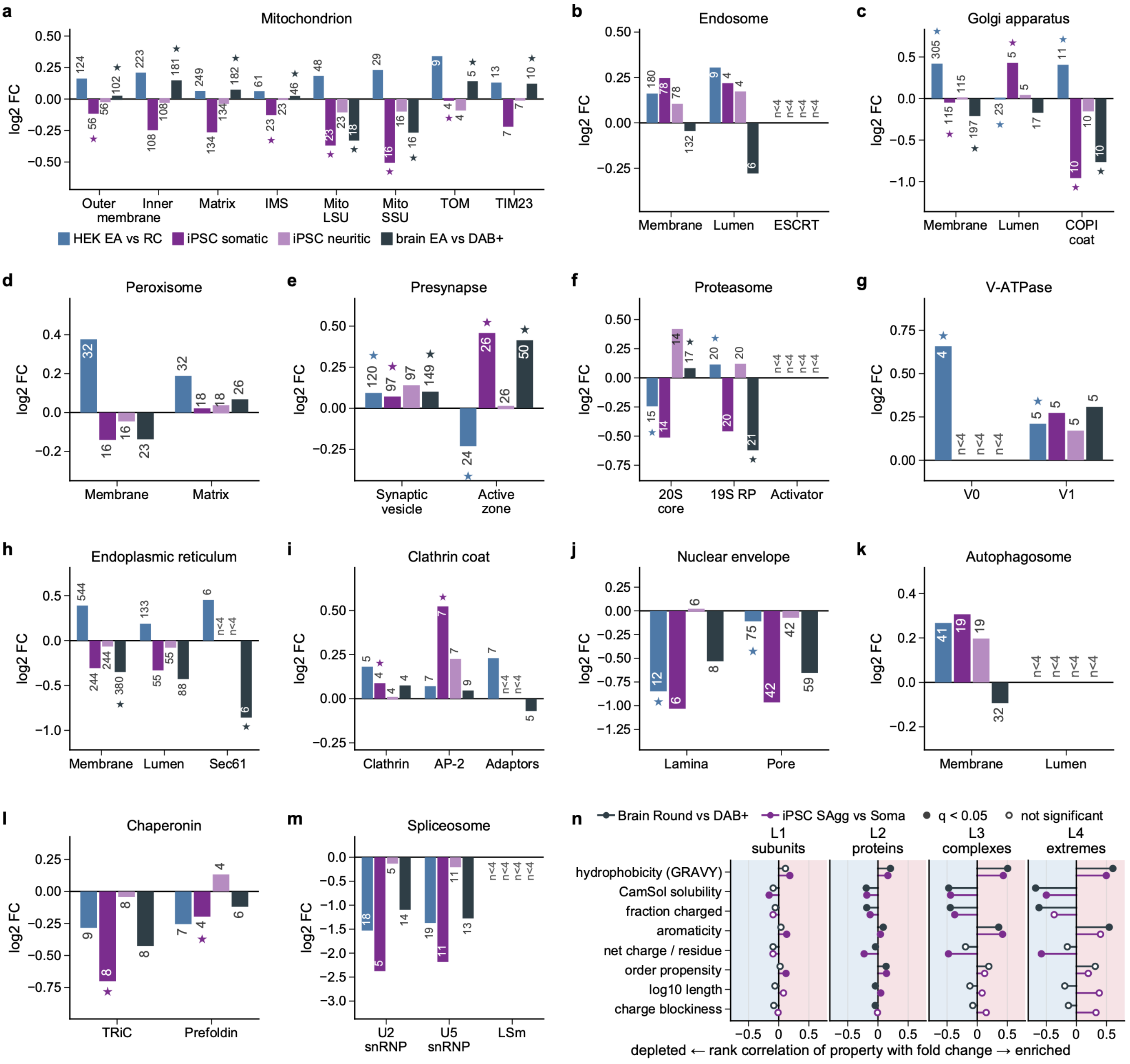


***Figure S8. Submodule structure of organelles and complexes in the aggregate, and sequence-property correlations across scales.***
*(a–m) The four bars are, left to right, HEK EA versus RC (−BafA1), iPSC somatic aggregate versus soma, iPSC neuritic aggregate versus neurites, and brain round aggregate versus DAB+ neuron; key in (a). A module score is the mean over its members after each protein is centred within the contrast; each bar is the group difference of those scores (brain, the mean of the per-donor differences). Numbers are member counts; n < 4 marks too few members to test, with no bar drawn. Modules are GO Cellular Component terms made disjoint within each organelle, the more specific one keeping any shared protein. A star marks a module in at least one pair separating at FDR < 0.05 by at least 0.3 log2 in that contrast (limma moderated t on module scores, in brain on one aggregate-minus-host difference per donor, BH within contrast). One module exceeding another means a larger aggregate-minus-host difference, so in a depleted organelle it means less depleted. The y axis differs between panels. (n) Spearman correlation between each sequence property and the aggregate-versus-host log2 fold change in brain (dark grey) and iPSC neurons (purple), at four levels: within each complex with every complex its own control (L1, mean of per-complex correlations, n ≥ 4 subunits); across all quantified proteins (L2); across complexes, mean property against median fold change (L3); and within the top and bottom quintile of complexes (L4). Positive means a higher value goes with enrichment, so the negative CamSol correlation means the less soluble proteins are the enriched ones. Filled markers are BH q < 0.05 from 10,000 permutations. Almost every L2 marker is filled because n is large there; read position, not fill.*

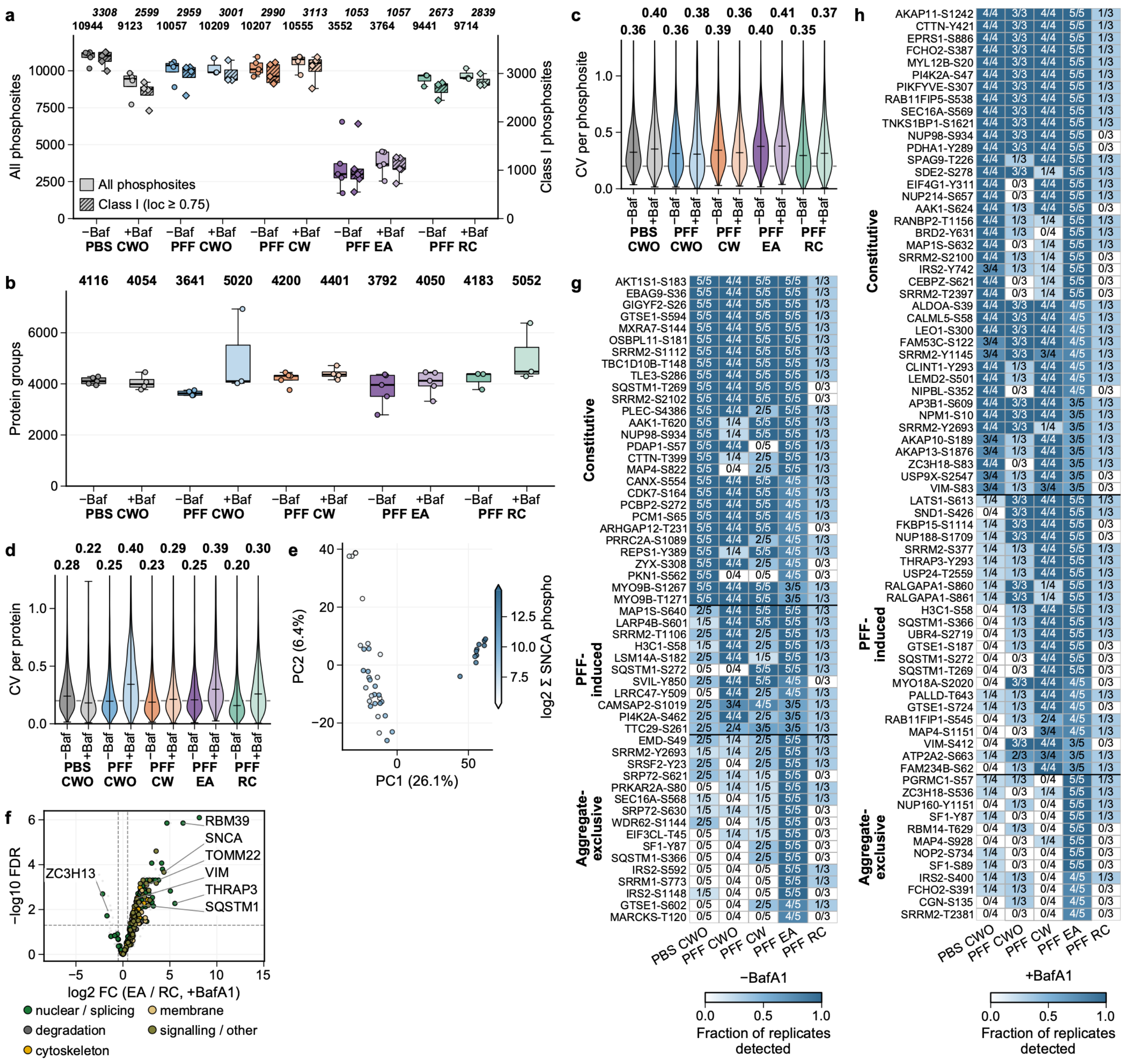


***Figure S9 Phosphoproteome data quality, α-synuclein phosphorylation load, and phosphosites partitioning into the aggregate.***

*(a) Phosphosites per sample by condition, all sites (left axis) and the Class-I subset (right axis). (b) Protein groups per sample by condition. (c, d) Coefficient of variation across replicates within each condition, per phosphosite on raw phosphopeptide intensities (c) and per protein on proteome intensities (d); means above each violin. (e) PCA of the Class-I phosphosite matrix (41 samples), coloured by summed α-synuclein phosphosite signal (seven detected SNCA sites). Excised aggregates separate along PC1 and carry the highest signal, aggregate-free cells the lowest, so the dominant axis of phosphoproteome variation corresponds to α-synuclein phosphorylation load. (f) Phosphosite log2 fold change, excised aggregates versus residual cell body under BafA1 (limma moderated t, each site normalised to its parent protein; dashed lines, |log2 FC| > 0.5 and BH-FDR < 0.05), the companion to the uninhibited comparison in Figure 6c. Coloured, sites of the genes in five hand-curated functional groups; grey, all other sites. (g, h) Phosphosites detected in excised aggregates (≥ 3 of 5 replicates) and absent from the matched residual cell body (≤ 1 of 3), without (g) and with (h) BafA1. These carry no quantifiable host signal and so cannot enter the comparison in (f) or Figure 6c. Cells give replicates detected (k of n), shaded by that fraction. Sites are grouped by where the mark exists outside the aggregate: constitutive, already present in fibril-naive cells; PFF-induced, still detectable outside; and aggregate-exclusive.*

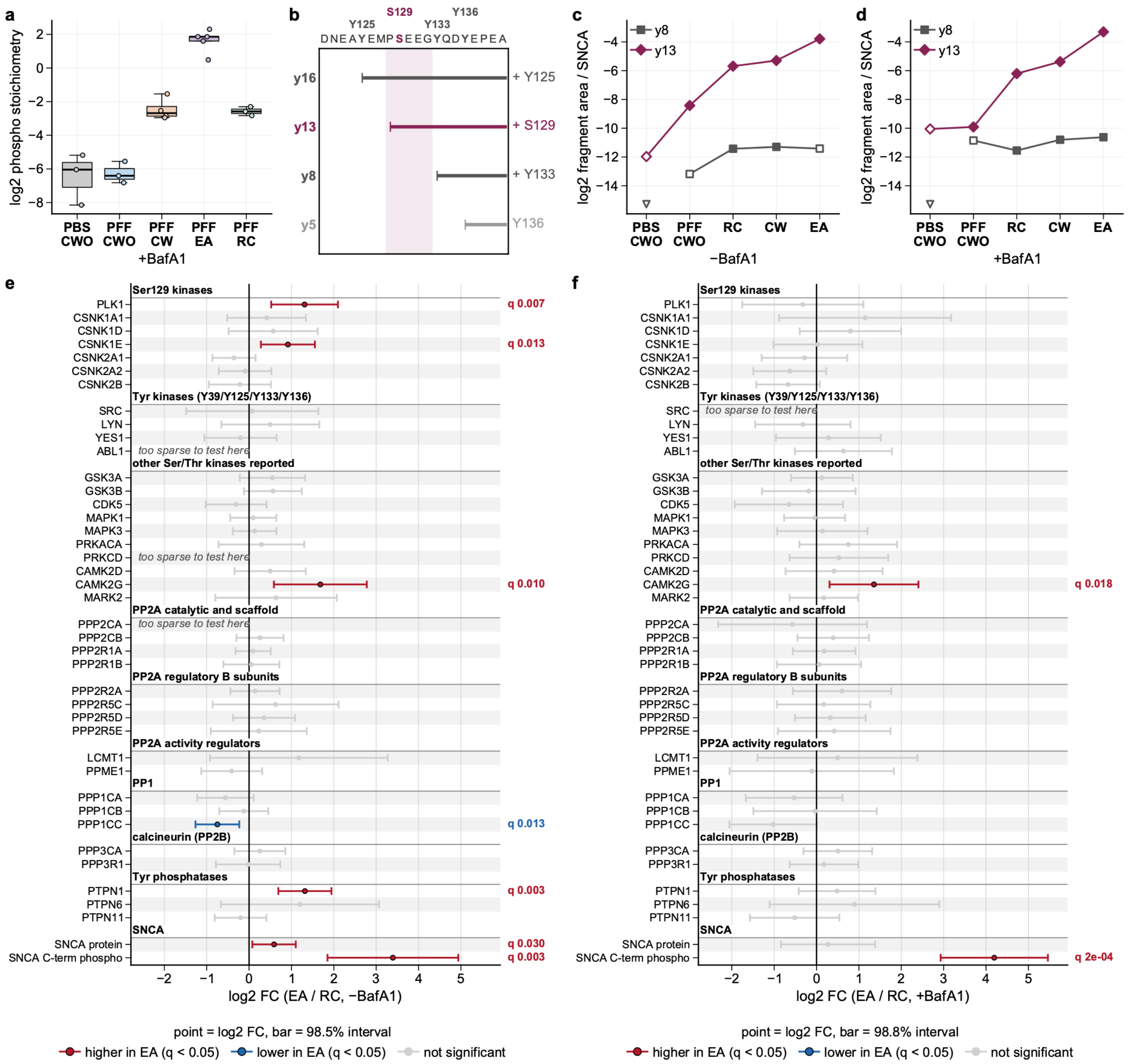

***Figure S10. Stoichiometry, site localisation and candidate enzymes for α-synuclein C-terminal phosphorylation.***
*(a) All phosphorylated precursors of the SNCA C-terminal tryptic peptide (residues 103–140, any localisation) under BafA1, proteome-normalised per condition (log2 phosphopeptide minus mean SNCA protein). Points are wells. (b) Coverage of the same peptide (121–140 shown). Each bar is the residues covered by one y ion, so the acceptors an ion can carry are read as a difference between ions. The shaded window (128–132) is the stretch gained between y8 and y13 and holds Ser129 as its only acceptor. (c, d) Summed peak area of the phosphorylated y8 and y13 ions across the dissection series without (c) and with (d) BafA1, without run normalisation and divided by the condition mean of SNCA protein; each point is the log2 of the mean over all wells of the condition, with an undetected well counted as zero. Open symbols, ion not detected in every replicate; symbols on the axis floor, never detected. Absolute intensities are not comparable between ions, so only the change across fractions is interpretable. (e, f) Protein-level log2 fold change, excised aggregates versus residual cell body (EA / RC), for every kinase and phosphatase with a reported link to α-synuclein, without (e) and with (f) BafA1. Bars are intervals at the level matching the Benjamini-Hochberg threshold across all tested rows, so the nominal level differs slightly between arms. Red and blue, q < 0.05; grey, not significant. Omitted are twelve with no protein group in any fraction (PLK2, PLK4, GRK3, GRK5, LRRK2, LCK, HCK, FGR, PTK2B, PPP2R2B, PPP2R2C, PPP2R5B) and sixteen detected in too few replicates to test (PLK3, GRK2, GRK6, FYN, ABL2, SYK, PRKCA, PRKCB, DYRK1A, PPP2R2D, PPP2R5A, PTPA, PPP3CB, PPP3CC, PTPN12, PTPRA). SNCA protein and its C-terminal phosphopeptide are shown for scale.*
